# Crystal structure of *E. coli* Nissle 1917 flagellin reveals novel features that modulate bacterial motility but not TLR5 recognition

**DOI:** 10.64898/2026.03.27.714778

**Authors:** Johanna Jakob, Michael B. Braun, Katharina Hipp, Iris Koch, Gaopeng Li, Pascal Felgner, Maria Jose Giralt Zuniga, Hannah Raasch, Corinna Ghering-Khav, Andras Szolek, Abdelhakim Boudrioua, Thomas Hagemann, Samuel Wagner, Thilo Stehle, Liudmila Andreeva, Marc Erhardt, Michael Hensel, Julia-Stefanie Frick, Alexander N. R. Weber

## Abstract

The probiotic *E. coli* Nissle 1917 (EcN) strain is known to promote intestinal homeostasis via flagellin, the protomer of its motility apparatus, the flagellum. The flagellin of EcN shows atypical features, namely a hypervariable region (HVR), whose structure and significance have remained elusive. We therefore determined the crystal structure of the *E. coli* Nissle 1917 flagellin FliC at a resolution of 1.2 Å which revealed an unusual domain architecture: the canonical D1 domain was found connected by an extended linker to an extensive HVR whose D2, D3 and D4 domains form an outer domain (OD) which surrounds the filament core comprised of conserved domains D0-D1. Using both recombinant proteins and gene-edited EcN strains expressing mutant flagellins, the functional requirement for these unique features was subsequently studied for effects on immune recognition on intestinal epithelial and immune cells, as well as on flagellar protein expression, assembly and bacterial motility. While human and mouse TLR5 immune recognition of flagellar proteins or intact bacteria was only moderately affected by removal of linker or D4, especially linker removal reduced protein stability and bacterial motility in both soft agar and liquid media swimming assays. Interestingly, depending on the environment, D4 or HVR removal had different effects on motility and surface structure. Finally, a site-directed mutagenesis approach highlighted that loss of TLR5 recognition strictly entails loss of motility but not vice versa. Our data indicate that specific HVRs/OD might be relevant for motility of *E. coli* Nissle 1917 in specialized environments, but not for immune recognition. Moreover, we find mutational tolerance is greater for immune recognition than for motility, providing new insights into bacterial adaptation to the host environment.

## Introduction

Flagella-mediated motility is a common and essential fitness trait in many microbes and enables bacteria to access nutrients or certain niches, and form biofilms while avoiding toxicity [1]. The complex bacterial flagellar apparatus hinges on the protein flagellin as its most abundant building block or protomer [2]. In *Escherichia coli*, an estimated 20,000-30,000 subunits of *fliC*-encoded flagellin make up each of the several 10 µm-long and 0.02 µm-wide flagella [3]. The inner core of this assembly comprises two sheaths formed by the D0 and D1 domains of flagellin. Central and outer regions of the flagellum are formed by consecutive additional flagellin domains, e.g., D2, D3 and D4, originating from a single polypeptide that folds back on itself in an antiparallel fashion, i.e., N- and C-termini join in the D0 domain as two antiparallel alpha helices. Whereas D0 and D1 are highly conserved across bacterial species, the other domains vary in number and residue length and are thus referred to as ‘hypervariable region’ (HVR). Especially regions beyond the D3 domain show great sequence and length variability: For example, *Salmonella enterica ssp. enterica* serovar Typhimurium (ST) FliC contains only domains D2 and D3 [4, 5], whereas *E. coli* Nissle 1917 (EcN), a well-known probiotic strain of *E. coli,* was found to contain a long HVR of 326 amino acids with unknown structure [6]. Even for different E. coli strains, flagellin sequence varies greatly (Fig. S1), with sequence identity in the HVR dropping as low as 29% (Fig. S2).

In a vertebrate host, flagellin also influences bacterial immunogenicity, serving as a microbe-associated molecular pattern (MAMP) sensed by the innate immune system via a pattern recognition receptor (PRR). Specifically, residues in the D1 domain are sensed by the PRR Toll-like receptor 5 (TLR5) [7]. Based on the crystal structure of FliC from *Salmonella enterica ssp. enterica* serovar Dublin in complex with zebrafish TLR5 [8] and synthetic biology studies [9], the FliC α-helices in the D1 region bind to TLR5 laterally, in the primary interface between the leucine-rich repeat (LRR)-NT and LRR10 regions, with LRR9 protruding out as a loop thereby forming a binding cavity for FliC. A second interface involves binding of the C-terminal αND1b to LRR12/13 of a second TLR5 molecule stabilizing homodimer formation of two TLR5:flagellin complexes in a 2:2 quaternary complex [8]. For signaling, a third binding interface is required that involves the D0 domain [10]. However, this seems to be unimportant for binding. Recent structural comparisons of fish and mammalian TLR5 indicate that this complex is likely an inhibitory arrangement [11, 12], so that the structure of an active ligand-receptor complex for the mammalian system has yet to be resolved. Mutagenesis studies indicate that residues 89-96, especially R90, and I411 within the *Salmonella* FliC D1 domain interact with A268 in human TLR5 [10, 13]. Binding of TLR5, which is expressed mainly on the basolateral sides of the intestinal epithelium, leads to mitogen-activated protein kinase and MyD88-dependent activation of NF-κB, which culminates in the secretion of pro-inflammatory cytokines (e.g., IL-6 and IL-8) and antimicrobial peptides (e.g., human beta defensin 2, hBD2) [14–16]. Mucosal immune cells also bear TLR5 and may respond to flagellin in case barrier integrity is compromised [17]. This is the case in inflammatory bowel disease such as Crohn’s disease [18, 19]. Recent work from our laboratories showed that flagellin from probiotic bacteria like EcN may have a regulatory effect on intestinal inflammation, specifically in a DSS-mediated experimental colitis mouse model [6]. We observed that EcN, but not other commensal *E. coli* strains, ameliorates DSS colitis in a flagellin- and IL-22-dependent manner. Paradoxically, these effects required TLR5 in conventional dendritic cells and yet involved the HVR, i.e. a region outside the classical TLR5 recognition motif [6]. Deletion of flagellin or parts of the HVR caused a loss of the probiotic effects. This begged the question whether structural features of the EcN flagellin might modulate TLR5 recognition and other aspects of EcN fitness, e.g., motility, a key property of bacteria navigating the intestinal space, including the mucus layer separating intestinal content from the (TLR5-expressing) intestinal barrier. The atomic structure of EcN FliC has not been determined, but recent structural studies and its length suggest it might adopt a fold identified in certain pathogenic and soil bacteria recently termed ‘outer domain sheath’ (ODS), including D2, D3 and an additional D4 domain [20].

Thus, although flagellin structure seems to have a profound effect on immune modulation and motility, the structure-activity relationships are not well understood, especially when considering potential “trade-offs” between immune recognition and motility. Sequence variation that would enable immune evasion may be counteracted by the need to retain motility; thus, changes at TLR5 binding interfaces are balanced against strong intradomain (e.g. between alpha-helices in D0 and D1) as well as homodimer contacts to guarantee flagellin folding and flagellar assembly [21]. One study recently investigated the effects of mutations in ST FliC on recognition by human and mouse TLR5 [22]. Interestingly, in an *in vitro* cell activation assay, recognition by human but not mouse TLR5 could be completely evaded by a variant of ST FliC carrying a single point mutation (R90A). In comparison, strains expressing ST FliC R90A retained 80% bacterial motility. In general, human TLR5 recognition was more sensitive to mutation compared to motility, whereas mouse TLR5 recognition was more robust and motility was lost more readily. In mice, but not in humans, *Salmonella* would therefore struggle to avoid TLR5 recognition while retaining motility. Another recent study focusing on human TLR5 found that a high proportion of flagellins in the human microbiota, e.g., *Roseburia hominis* FlaB, are ‘silent’ because they bind to TLR5 but fail to activate downstream signaling [23]. Engraftment of ST FliC D0 restored TLR5 activation in purified *Roseburia* FlaB *in vitro*, highlighting the contribution of this domain to TLR5 activation [10, 23]. However, the stability and ability of these hybrid flagellins to form flagella and the resulting effects on motility were not assessed. Thus, it is unknown whether the ‘silent’ D0 domain of *Roseburia hominis* FlaB allows for motility in other flagellated bacteria, such as *E. coli* or ST.

To gain a deeper understanding of the structural basis for the probiotic effect of the EcN flagellin and the relationships between flagellin structure, TLR5 recognition and motility, we first solved the crystal structure of EcN FliC and discovered that the HVR sequence adopts another well resolved domain, D4. Moreover, and unexpectedly, D1 and D2 were separated by an extended antiparallel β-sheet linker. We examined the effects of these unique structural features on immune recognition by human and mouse TLR5 as well as motility. This analysis showed that both unique features are redundant for TLR5 recognition and secretion of flagellin from bacteria. However, both linker and D4 domain emerged as essential for optimal bacterial motility in certain environments. EcN D4 appeared to represent a novel OD. Although deletion of the linker and the D4 compromised motility, a complete removal of the HVR was, surprisingly, tolerated, but depending on context. Our data indicate that under *in vitro* conditions, ODSs are not essential for motility *per se*, but our study highlights the unique D1-D2 linker in the flagellar EcN core as a critical context-dependent determinant of bacterial motility. Moreover, using mutagenesis studies we found that loss of TLR5 recognition strictly entails loss of motility but not vice versa, providing new insights into host-bacterial adaptation.

## Results

### The crystal structure of the atypical EcN flagellin contains an extensive ODS with a rigid linker

To gain insight into the structure of EcN flagellin, especially its HVR, we expressed different constructs of the entire EcN FliC in *E. coli* BL3 cells as His-tagged fusion proteins (see Methods, Fig. S3 and Fig. S4) for subsequent crystallographic analysis. Whereas full length and a D0 deletion construct (ΔD0, Δ1-47, Δ555-595) did not yield crystals, a construct with additional deletion of D1 (ΔD0+D1, Δ1-175, Δ499-595) could be crystallized and diffracted. As molecular replacements (ST FliC PDB 1IO1), as well as native and seleno-methionine phasing did not yield results, heavy atom (Se-Urea) soaking was performed, allowing for solving the native structure at 1.65 Å resolution. As the D1 is critical for TLR5 interaction, another attempt to gain D1 information was made using additional shortened constructs, FliC D1+2 (Δ1-61, Δ208-451, Δ542-595) and FliC D12(Δ1-61, Δ208-451<S, Δ555-595), which resulted in crystals that diffracted to a resolution of 2.03 Å and 1.75 Å. Since the D2 domain was present in both sets of constructs, it was superimposed with minimal differences (RMSD 0.4 Å over 275 aa), so that a combined model of the D1 to D4 domain was built (Fig. 1A). The structure showed several typical features: firstly, a helical D1 domain with anti-parallel coiled-coil structure that aligned well with the ST FliC (PDB 1IO1, Fig. S4E, RMSD 0.57 Å); secondly, a globular D2 with a three stranded antiparallel β-sheet and one α-helix forming an αβββ-sandwich; and thirdly, a globular D3 domain consisting of two αββ-sandwiches where the four β-strands form a β-sheet with the two α-helices laying on the same side. Surprisingly, another domain, D4, with a similar αββ-sandwich topology but 90° rotated β-sheet orientation (Fig. 1B), was discovered. The tip of D4 showed an unusual fold, where three strands form an antiparallel triangle, with β-sheet-like interactions, termed here β-triangle. Thus, the HVR sequence previously described [6] contributed to a canonical D3 and to a unique D4 domain. Compared to ST FliC, the positioning of D1-D3 deviated strongly. This was primarily due to a linker separating D1 from D2, resulting in minimal contacts between the two domains (Fig. 1C). Instead, in ST FliC, the transition between D1 and D2 is seamless and generates numerous interactions between the D1 and D2 domain. The length of the linker is about 30 Å, with the first 21 Å, from D1 to D2, stabilized by ten hydrogen bonds and one ionic interaction between Q500 and K505. In the second, 9 Å-long part of the linker, the N-terminal strand forms two hydrogen bonds with the C-terminal strand, which itself forms a β-hairpin. This C-terminal β-hairpin also interacts with the D2 core fold, which additionally stabilizes the linker region. Three structures of the FliC D12 construct were solved from two crystals (one asymmetric unit contained two protomers). This allowed for a structure alignment that helped to visualize the differences in the relative alternative positions of D2 (Fig. 1D). The angle between the center of mass (COM) of D1, the Cα of P507 at the tip of D1 and the COM of D2 varies between 130° and 136°. The angle between the Cα of P507, the COM of D2 and the carbonyl C of T206, a point close to the transition to D3, varies between 92° and 107°. Thus, the unusual linker would provide a certain degree of flexibility between D1 and subsequent domains. We were intrigued to explore how these unique features of the EcN FliC D1-D4 might affect flagellar structure and modelled on the published cryo-EM structure of *Bacillus subtilis* flagellum (PDB 5WJT) using the D1 for alignment. The two resulting models differ in their shape as a result of the different HVR domain orientations. The model with the HVRs closer to the D1 resulted in a closed flagellum of 211 Å diameter, where the HVRs form an outer ring-like structure. In the other model with 252 Å diameter, D3 and D4 are moved outwards, opening the gap between inner constant domain superstructure and the outer HVR superstructure (Fig. 1E). In these predicted filaments most interactions would occur between D0 and D1 domains. Between the two plausible models of the flagellar assembly, which include the HVR, we noted a difference in the connectivity (Fig. 1F). In the more ‘closed’ model, the flagellum would be narrower (211 Å) and HVR domains are closer to the constant domains, with more putative interactions between different HVR domains. The more ‘open’ model shows less contact between the domains but yields a wider flagellum (252 Å). Collectively, our structural studies show EcN FliC adopts a flagellar fold with unique features, most importantly the D1-D2 linker and the outer D4 domain, which are compatible with the formation of a functional flagellum.

**Figure 1:**
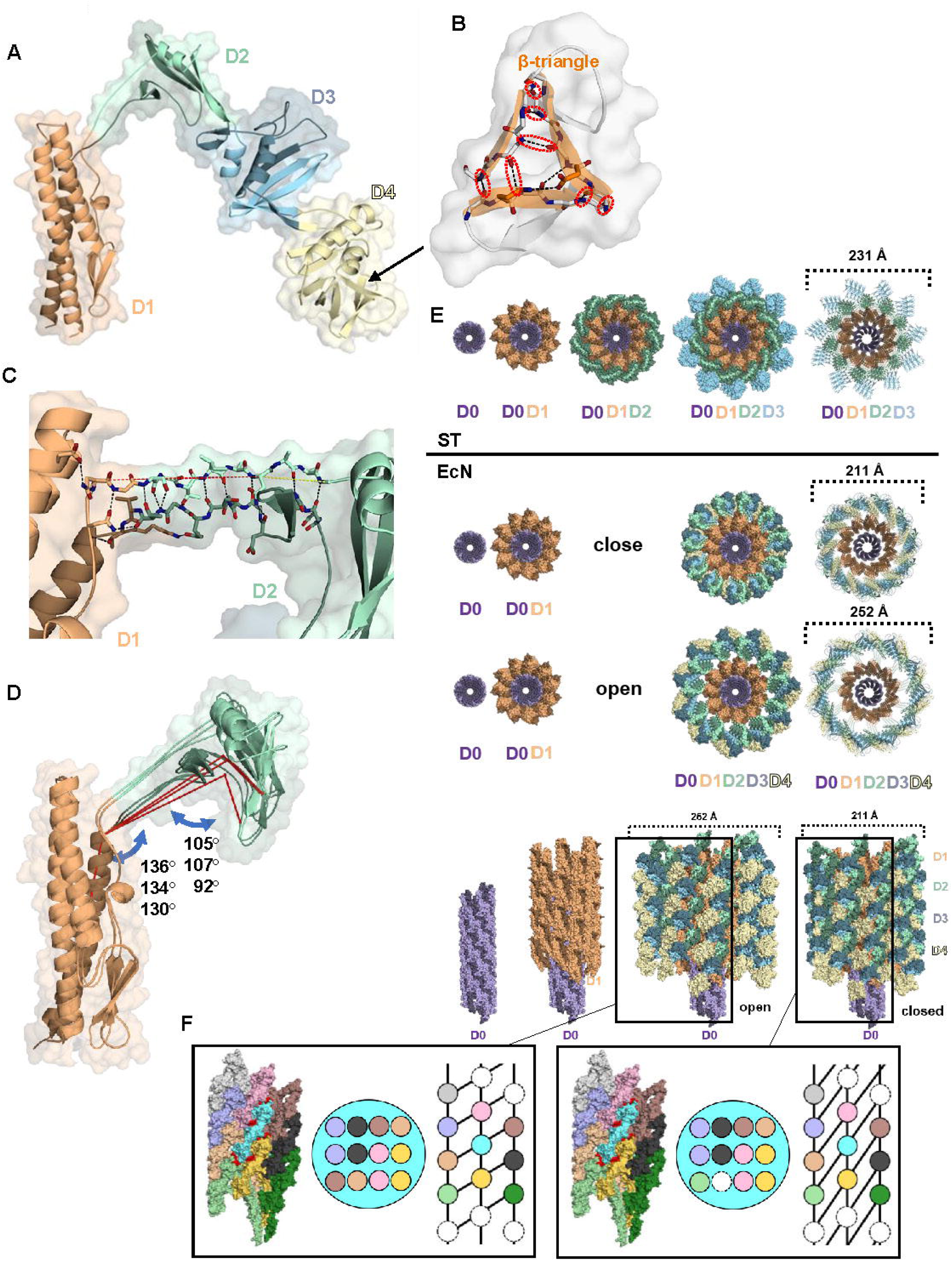
Structure of the EcN flagellin FliC shows a distinct D4 domain and D1-D2 linker. **(A)** Cartoon and surface representations of the EcN FliC model comprising the D1 (orange), D2 (green), D3 (blue) and D4 (yellow). **(B)** Side-view of the D4 β-triangle. **(C)** Interaction of the two strands of the D1-D2 linker. The hydrogen bonds and ionic interaction are depicted with black dotted lines. The distance between the D1 and D2 is depicted with a red dotted line (21 Å) and a yellow dotted line (9 Å). The region close to D2 is stabilized by a β-hairpin. Three different flagellum models. **(D)** Structure alignment of three EcN D12 structures, side view with angles between center of mass (COM) of D1, Cα of P507, the COM of D2, and carbonyl-C of T206, connected with a red line. The structures are shown in cartoon and surface representations, with D1 (brown) and D2 (green). **(E)** Comparison of ST and EcN FliC in flagellar context. Cartoon or surface representation axially viewed from the tip towards the stator of the flagellums, with D0 (purple), D1 (orange), D2 (green), D3 (blue) and D4 (yellow). Top: ST. Middle: closed model of the EcN flagellum. Bottom: open model of the EcN flagellum.

### Linker and D4 domain are dispensable for TLR5 recognition

A *Salmonella* Dublin FliC:zebrafish TLR5 complex [8] has long been used as a blueprint for flagellin recognition by human TLR5, although it has been proposed it may represent an inhibitory complex (discussed in [12]). Nevertheless, to gain an insight into structural aspects of EcN FliC interaction with TLR5, the EcN FliC and ST FliC structures were superimposed (Fig. S5). This showed that the EcN and ST D1 domains interacted well with TLR5. However, the EcN D2-D4 domains hardly interacted with TLR5. Of note, for the ST FliC, an agonist of both human, mouse and zebrafish TLR5 [11], its D2-D3 were predicted to clash with the second TLR5 molecule, providing further evidence that the described *S.* Dublin:TLR5 complex is unlikely to represent an active complex (Fig. S5).

We next sought to directly explore the relevance of the unique EcN structural features for TLR5 recognition, using HEK293T cells stably expressing human or murine TLR5. These were either challenged with purified flagellins, including the D0-D4 domains (full FliC), ΔD0, ΔD4, Δlinker or ΔHVR, or exposed to EcN bacteria in which the chromosomal *fliC* had been edited accordingly (see Methods). Purified ST FliC and/or aflagellated EcN (Δ*fliC*) were used as controls, respectively. In line with earlier results, the D0 seemed strictly required for both human and mouse TLR5 recognition. Interestingly, whereas the D4 was largely dispensable for recognition of purified flagellins at equimolar concentration, deletion of the linker slightly reduced IL-8 output (Fig. 2A, 2B). When bacteria expressing the corresponding FliC variants were used to stimulate hTLR5 and mTLR5 HEK29eT reporter cells, no differences were apparent for hTLR5 (Fig. 2C), but mTLR5 cells showed a slight reduction for ΔD4- or Δlinker-FliC-expressing bacteria (Fig. 2D). Thus the unique structural features of EcN FliC only moderately affect direct TLR5 recognition. As an alternative to HEK293T, we performed similar experiments in more physiologically relevant intestinal epithelial systems, namely, C2BBe1, an intestinal epithelial cell (IEC) line, and intestinal human organoid cultures, as well as immune cells, namely, the human macrophage-like THP-1 cell line and primary murine bone-marrow-derived dendritic cells (BMDCs). Intestinal C2BBe1 cells responded to purified full-length FliC and ΔD4 FliC with robust IL-8 release, but Δlinker, ΔHVR and ΔD0 FliC did not trigger release significantly above background (Fig. S6A). Human beta defensin 1 (hBD1) mRNA was induced comparably little between flagellin preparations (Fig. S6B), but hBD2 was strongly induced by all flagellins, with significantly more hBD2 mRNA induced by the Δlinker flagellin version, and significantly lower by ΔD0 flagellin (Fig. S6C). In similar experiments using whole bacteria, we noted comparable IL-8 release from C2BBe1 cells in response to all but Δ*fliC* (aflagellated) and *fliC* Δ*TLR5rs* bacteria (which lack the TLR5 recognition epitope), suggesting a TLR5-dependent response. Of note, mouse commensal *E. coli* strains, namely *E. coli* mpk, and *E. coli* MG1655 also triggered significantly lower IL-8 release (Fig. 3A), in line with earlier data [6]. HBD1 mRNA was not induced significantly (Fig. S7A), but the strong hBD2 mRNA induction (>1000-fold) followed IL-8 results in that *fliC* mutant strains Δ*TLR5rs* and Δ*fliC*, as well as *E. coli* mpk and MG1655 hardly induced any hBD2 mRNA (Fig. 3B). Interestingly, in this assay ΔD4 FliC-expressing bacteria also did not trigger hBD2 mRNA transcription (Fig. 3D). A preliminary test (n=1) in human enteroids stimulated with purified flagellins showed comparable IL-8 release, except ΔHVR and ΔD0 flagellins, which induced significantly less IL-8 than WT EcN flagellin (Fig. S7B). Switching to immune cells, which, unlike intestinal epithelial cells, are also competent for the LPS receptor, TLR4, we focused on entire bacteria only, as even minor traces of LPS in the purified protein preps would be expected to confound the data in a poorly tractable manner. On the other hand, the analysis of whole bacteria enables the effects of flagellin to be assessed in the context of other MAMPs naturally present in the same bacteria, and hence provides a physiologically relevant context. In co-culture with THP-1 cells, different MOIs of the EcN variants expectedly released very high levels of IL-8 and TNF, intermediate levels of IL-6 and IL-10 and low levels of IFNβ and IL-1β, but there were no significant differences between WT EcN and deletion mutants, mostly likely due to LPS acting as the overriding MAMP (Fig. S8). Exposure of primary BMDCs (Fig. S9) to flagellin-expressing bacteria resulted in downregulation of TLR5 (Fig. S10A), no effect on CD40 (Fig. S10B) and upregulation of CD86 and MHC-II (Fig. S10C), although there were again no significant differences between strains. The same applied to cytokine release, which was as strongly induced by but did not vary between different flagellins (Fig. S10E-H). These data are exemplified for IL-8 in the heat map of Fig. 3C. Collectively, we can conclude that immune cells, due to the dominance of LPS, cannot distinguish different bacteria based on different forms of flagellin. However, in cells with strict TLR5 dominance (transgenic TLR5-HEK cells and epithelial cells), deletion of the linker had a subtle effect on recognition at the level of purified proteins (albeit not as great as ΔD0). In good agreement with the structural analysis (*cf.* Fig. 1E), neither the D1-D2 linker nor the D4 domain thus seemed to have a strong very effect on immune recognition by immune and epithelial cells.

**Figure 2:**
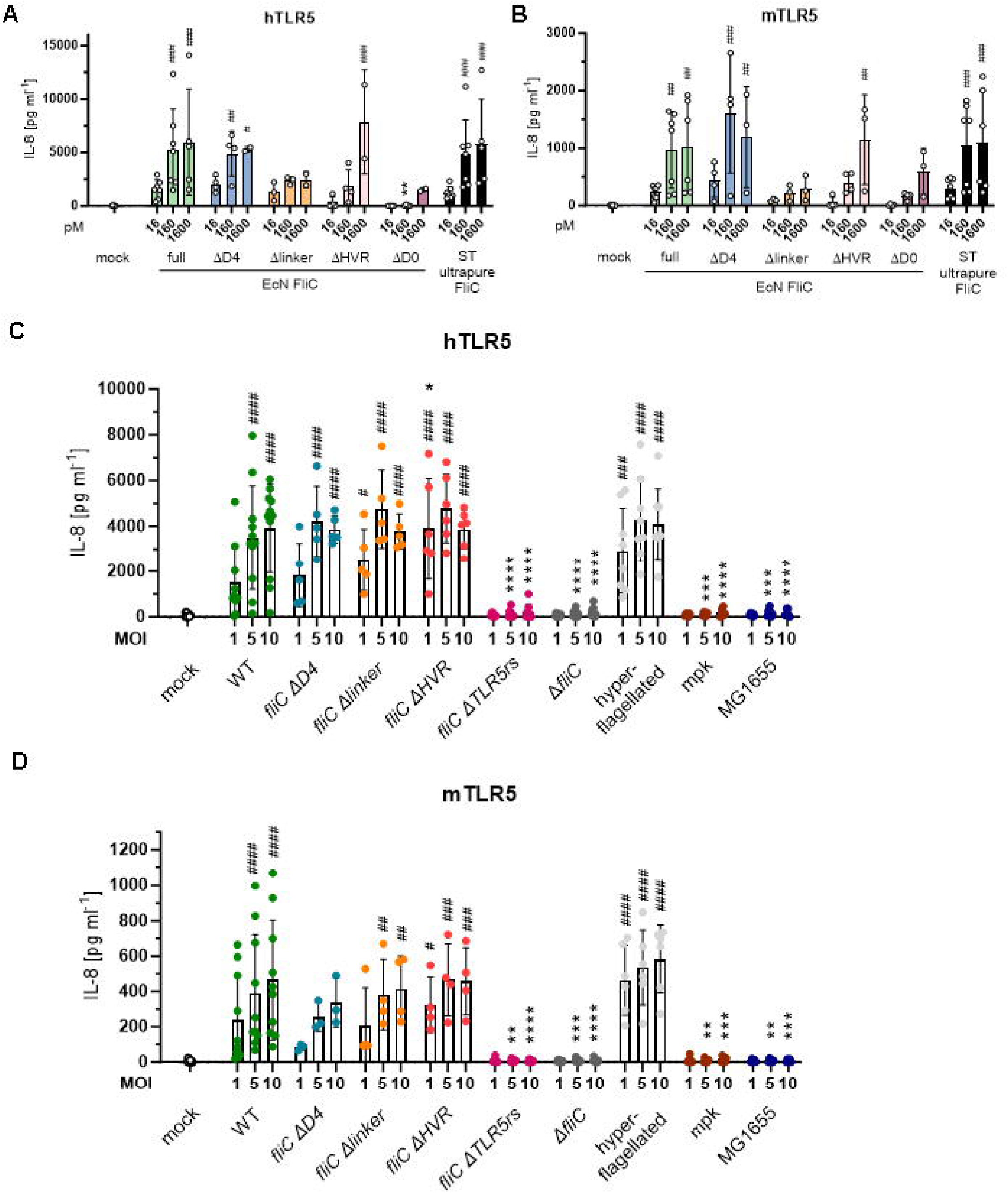
The D1-D2 linker affects TLR5 recognition of purified EcN flagellin, but not flagellated bacteria. IL-8 cytokine ELISA of **(A, C)** human (h)TLR5 or **(B, D)** murine (m)TLR5-expressing HEK293T cells stimulated with **(A, B)** recombinantly isolated FliC proteins or with **(C, D)** gentamycin-killed bacteria for 24 h. Pooled data, n ≥ 3 experiments; statistics: two independent tests were done: 2-way ANOVA with Dunnett correction sample versus mock indicated by #; 2-way ANOVA with Dunnett correction sample versus full-length FliC indicated by *

**Figure 3:**
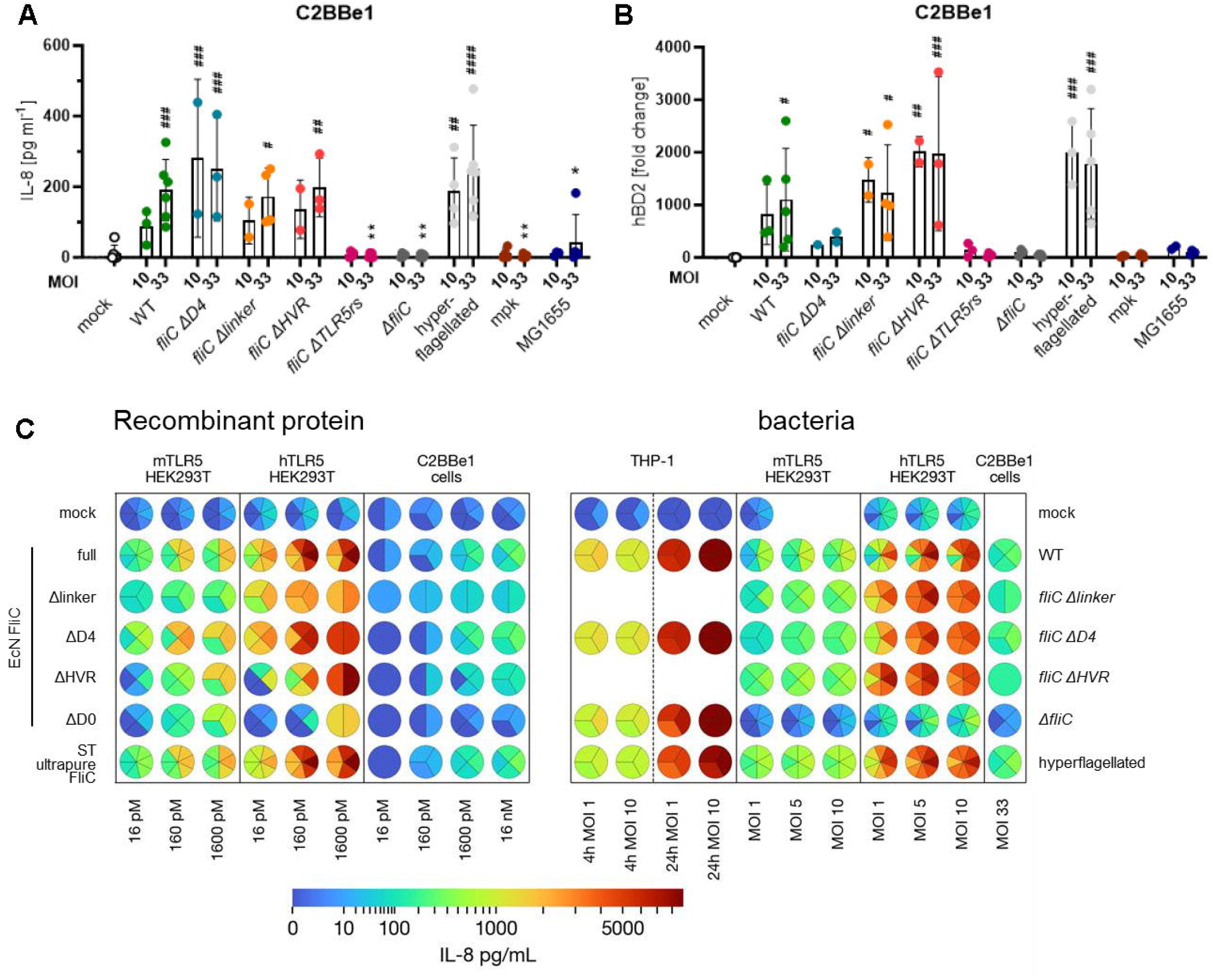
Subtle discrimination of purified EcN flagellins but not flagellated bacteria by intestinal epithelial and immune cells. **(A)** IL-8 cytokine ELISA and **(B)** human beta defensin 2 (hBD2) expression of C2BBe1 cells 6 h after stimulation with gentamicin-killed bacteria. Pooled data, n ≥ 3; statistics: two independent statistical tests were performed, 2-way ANOVA with Dunnett correction sample versus mock indicated by #; 2-way ANOVA with Dunnett correction sample versus EcN WT indicated by *. **(C)** Heat maps summarizing IL-8 cytokine ELISA data of THP-1, mTLR5-, hTLR5-HEK293T and/or C2BBe1 following stimulation with recombinantly isolated FliC proteins (left) and gentamicin-killed bacteria (right) after 24 h or 6 h (C2BBe1). Pooled data, fractions of pie charts indicate technical replicates (THP-1 data) or mean value of n independent experiments.

### Linker and D4 domain affect flagellar expression, assembly and motility

Apart from immune recognition, the previously reported probiotic effect of EcN might also be due to different swimming capabilities, e.g., different encroachment of the intestine. To explore the effect of the deletions outlined above on motility, we first investigated whether strains expressing the mutant flagellins chromosomally were able to secrete flagellin and form flagella by assessing flagellin release in the culture media and negative-stain transmission electron microscopy (TEM) micrographs of different EcN mutants. A preliminary analysis of culture supernatants by SDS-PAGE and silver staining showed that all deletion and mutation constructs were detectable and secreted to levels comparable with wild type (WT) EcN flagellin, except ΔHVR and ΔTLR5rs flagellin (Fig. S11A). To assess the stability of the proteins themselves, we conducted thermal stability assays with recombinant purified proteins, which showed that deletion of D4, and even more so the linker, reduced stability, whereas deletion of the entire HVR did not (Fig. S11B). To check if this had an effect on flagellation, transmission electron microscopy (TEM) of all EcN mutant strains was performed. All strains, including the *fliC* Δ*HVR*, showed flagella, apart from the Δ*fliC* (expected) and the *fliC* Δ*TLR5rs* variant (Fig. S12), indicating this deletion might not only abrogate the reported TLR5 recognition but also proper flagellar assembly. Next, the ability for motility was assessed in soft agar assays [24](Fig. S13). Interestingly, quantification of swimming radii (Fig. 4A; 4B) showed that *fliC* Δ*D*4 and *fliC* Δ*linker* deletions were associated with a significantly compromised swimming distance compared to EcN WT but retained residual motility relative to the non-motile aflagellated *fliC* Δ*TLR5rs* and Δ*fliC* strains. Surprisingly, a strain that lacked the entire HVR (*fliC* Δ*HVR*) was as motile as WT EcN, suggesting that D4 in its full EcN FliC context contributes to motility and its truncation partly destabilizes the flagellum, whereas full deletion of the HVR, i.e., deletion of T180-E500, can yield a functional flagellum, despite lacking the outer domains. Interestingly, single-cell motility analysis (Fig. 4C) confirmed an essential requirement for the linker in liquid environment. A deletion of the D4 domain did not affect average swimming speed, whereas deletion of the entire HVR significantly reduced swimming velocity in this assay. Thus, it appeared that different environments (viscous soft agar vs. liquid) may require certain flagellar features for optimal motility. One possibility is that different surface structures of the flagellum create different levels of traction. Indeed, analysis of individual flagella by TEM showed interesting differences in surface granularity, namely that, compared to WT the *fliC* Δ*HVR* mutant showed a both thinner and smoother flagellar filament (Fig. 4D, 4E), whereas the *fliC* Δ*linker* filament was only slightly thinner but retained a more granular surface appearance. Moreover, when analyzing the wave form of these flagella, *fliC* Δ*HVR* and *fliC* Δ*linker* flagella showed smaller distances between consecutive pitches (i.e. a smaller wave-pitch distance; Fig. 4F). We speculate that especially in liquid environments, a smoother surface makes motility less efficient, compared to movement in a more viscous soft agar. Moreover, the low motility of the *fliC* Δ*linker* mutant in both environments (*cf.* Fig. 4A/4B vs 4C/D) – despite a more WT surface granularity but lower wave pitch distance, like in the *fliC* Δ*HVR* mutant – may be explained by its significantly lower filament stability (*cf.* Fig. S11), which would be expected to compromise flagellar performance independent of the environment. Collectively, our analysis indicates that the HVR as a whole, the atypical linker and D4 domain do not hinder TLR5 recognition, but they affect bacterial motility differently depending on the environment by modulating protein stability, surface roughness and wave-form of the filament.

**Figure 4:**
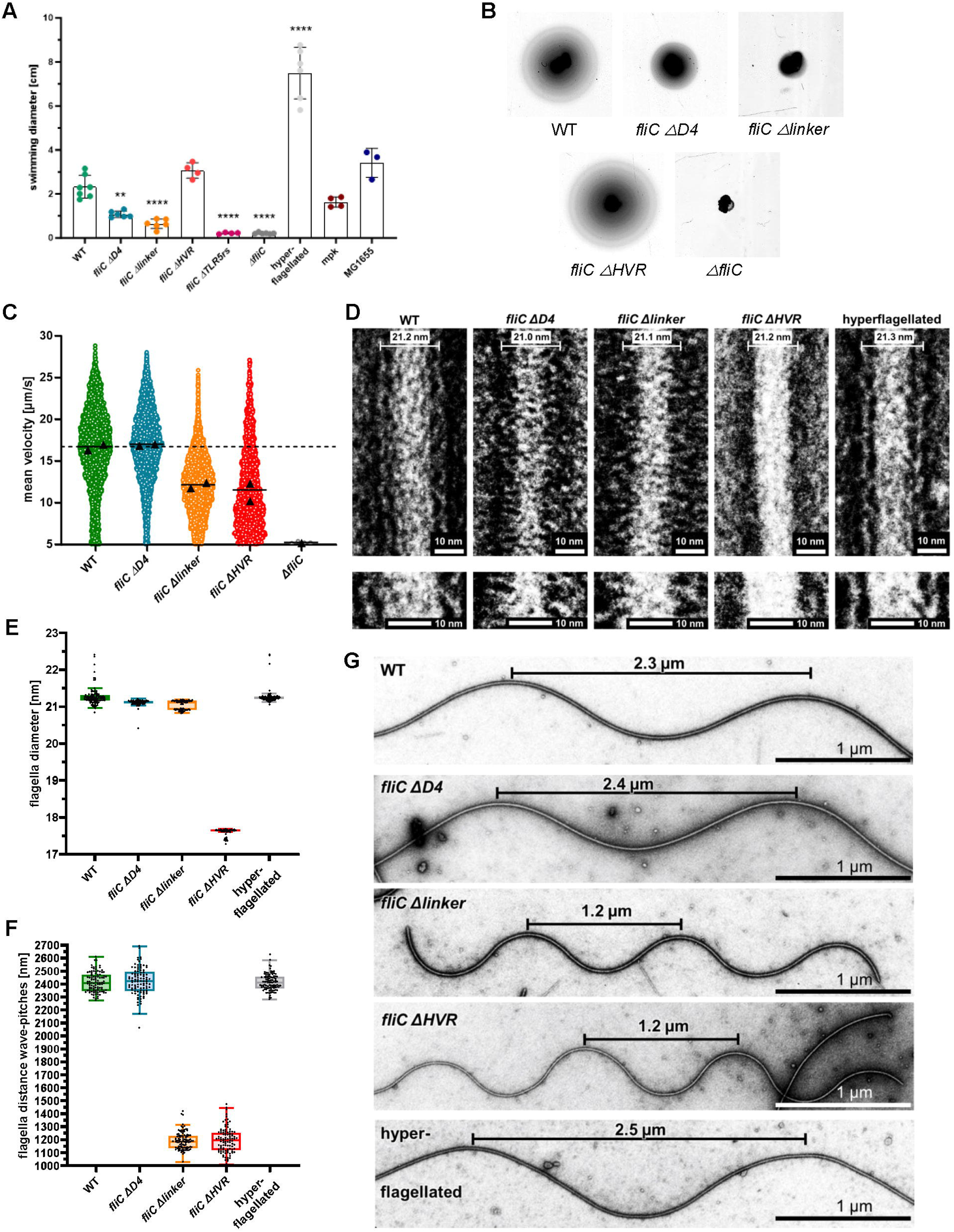
Linker and D4 domain affect motility. **(A)** Soft agar swimming distances of FliC mutants depicted as mean of measured diameters d, n ≥ 3; statistics: ordinary one-way ANOVA with Dunnett correction sample versus EcN WT indicated by *. **(B)** Representative images for selected bacteria. **(C)** Mean single-cell swimming velocities in liquid media of wild-type and mutant strains obtained from exponentially growing cultures. Each dot represents the mean velocity of a trajectory of an individual bacterium (≥3 s), with dots colored by replicate. At least 1521 individual trajectories were analyzed per strain and replicate. **(D)** Representative TEM images of negative-stained isolated flagella from wildtype (WT), hyperflagellated (hyperflag.), *fliC* Δ*D4*, *fliC* Δ*linker*, and *fliC* Δ*HVR* strains after background subtraction. All samples were prepared as described in Materials and Methods. **(E)** Average filament diameters were quantified using ImageJ and are indicated for each strain. The diameter of 100 flagella per variant was determined by calculating the mean value from three measurements performed along each filament. Scale bar: 10 nm. **(F)** The average wave-pitch distances were quantified from 100 flagella per strain using ImageJ. Scale bars as indicated. **(G)** Representative TEM images of isolated flagella showing distinct wave-patterns between the different variants. Redlines indicate the measured wave-pitch distances.

Thus far, our data indicate that TLR5 recognition of EcN flagellin was less structurally sensitive than bacterial motility, so that deletions in EcN flagellin that significantly affect or ‘trade in’ motility still did not evade full TLR5 recognition. Next, we sought to probe whether more subtle modifications, such as point mutations, might generate EcN variants, in which motility was retained but TLR5 recognition evaded, as shown for ST FliC [22]. In light of the observed differences in sensitivity between human and murine TLR5, we exposed HEK293T expressing either of these receptors to different EcN mutants, which were generated based on comparisons of immunostimulatory (EcN and ST FliC) and TLR5-evasive flagellin (*Campylobacter jejunii*) mutants at the proposed TLR5 interface [21]. Most single mutants still caused significant secretion of IL-8, e.g. R93K, E94T and R530Q, 24 h after infection (Fig. S14). R91T seemed to be the exception and emerged as crucial for recognition by human and even murine TLR5, as it strongly reduced IL-8 secretion. We also tested combinations of the previous mutations, but these only highlighted the dominance of R91T for TLR5 recognition. We were curious about the effect of these mutations on motility (Fig. S15 and Fig. 5A). Evidently, ‘tampering with’ R91 completely crippled EcN motility (Fig. 5A) in that we were unable to identify any single or multi-mutants with abrogated TLR5 recognition but intact motility amongst the analyzed R91-mutated variants. Summarizing and correlating relative (to WT) NF-κB activity of TLR5 HEK293T with motility data (Table S2 and Fig. 5B/5C) suggested that greatest achievable reduction in TLR5 activity, to (only) about 70%, coincided with a motility reduction to 80% (R530Q) or even ∼30% (R91T E94T). Any further reduction of TLR5 activity below 60% resulted in a concomitant drop in motility to 40% or less (red quadrant). Conversely, several mutants with less than 50% motility retained a relatively high TLR5 reactivity (orange quadrant). This shows that motility seems far more sensitive to mutations than TLR5 recognition is. Alternatively phrased, EcN undergoing FliC mutations would probably be more likely to lose motility before avoiding TLR5 recognition.

**Figure 5:**
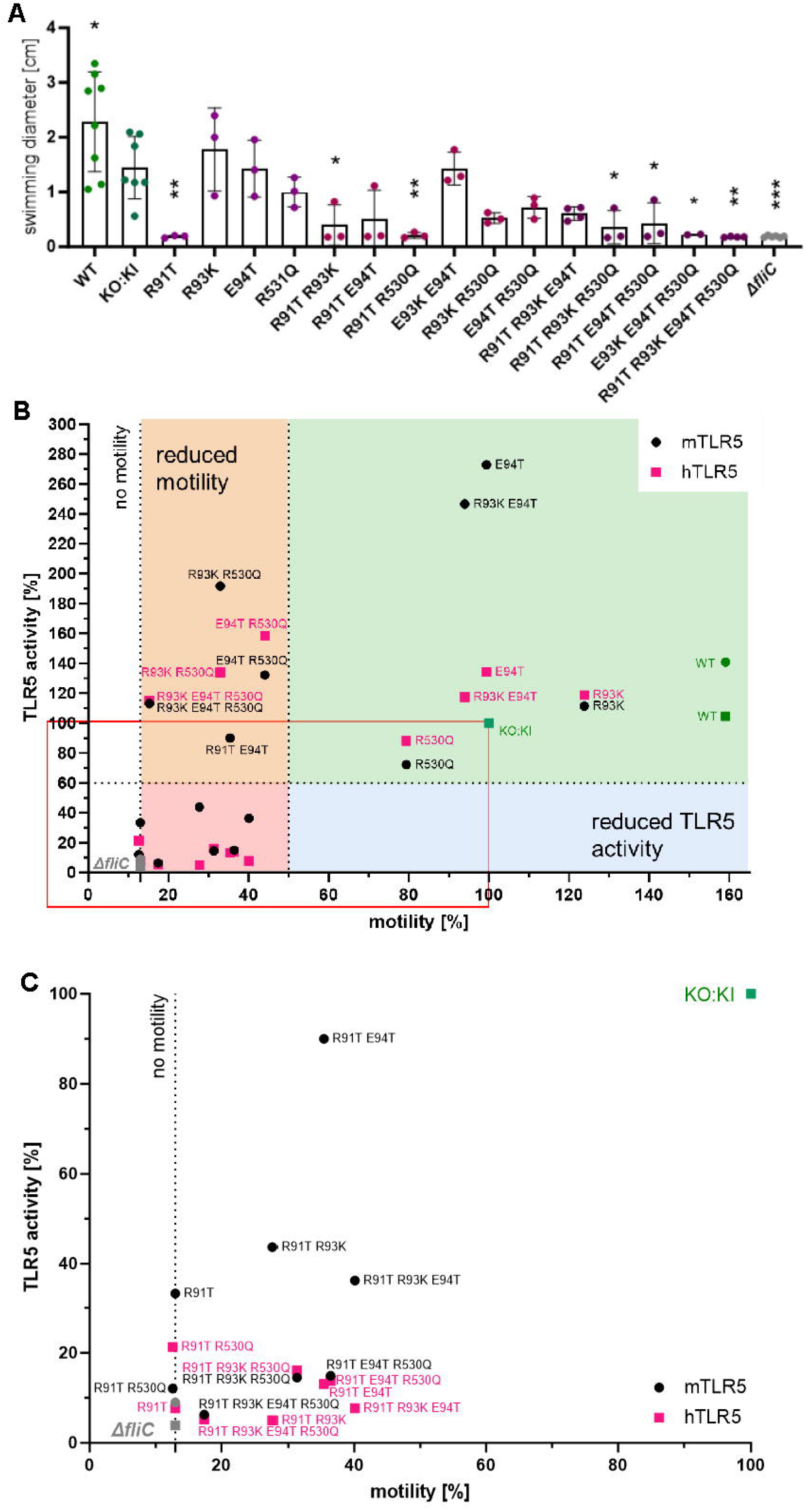
Point mutations in TLR5 interface impair motility but not TLR5 recognition. **(A)** Soft agar swimming distances of FliC mutants depicted as mean of measured diameters, n ≥ 3. Statistics: ordinary one-way ANOVA with Dunnett correction sample versus KO:KI indicated by *. **(B)** Graphical summary of normalized m- and hTLR5 activity plotted against normalized motility for each mutant. Dashed line on the left indicates lower limit for motility derived from bacterial volume on plate despite lacking motility.

## Discussion

EcN represents one of the most extensively studied probiotic *E. coli* strains. Although previous work implicated its flagellum, and especially its HVR, and TLR5 to be a critical mediator of probiotic effects in experimental colitis [6], these properties have not yet been linked to certain molecular or structural features. Here, we therefore determined the EcN HVR structure, highlighted unique features compared to the prototypical ST FliC, but probably not uncommon in *E. coli*. Most striking were the linker between D1 and D2 and the additional D4 domain.

Regarding the latter, our data about the existence of an additional OD is in good agreement with a recent cryo-EM study on flagellar filaments from enterohemorrhagic *Escherichia coli* O157:H7, enteropathogenic *E. coli* O127:H6, *Achromobacter*, and *Sinorhizobium meliloti* [20]. This study first described outer domain sheaths as a structural feature enabling cross-linking of flagellin protomers via D4 dimerization, thus allowing for prolonged *E. coli* tumbling. Moreover, it was speculated, albeit not proven, that a screw-like outer surface might provide additional traction in high viscosity environments such as the intestine. Our structural data revealed that probiotic EcN shares these structural features with pathogenic *E. coli* and that the observed D4 dimerization also applies to EcN flagellar structure. In fact, HVRs long enough to contain a D4 occur in many bacteria and the D4 sequence of EcN collects BLAST hits, e.g., in pathogens like *Shigella* spp (data not shown).

Interestingly, although this feature was not reported or functionally assessed before, the extended D1-D2 linker in EcN FliC is also present in atomic cryo-EM models of EHEC O157:H7 and EPEC O127:H6, two pathogenic *E. coli* strains. These structures show a linker of different amino acid sequences but similar length and angle to EcN (Fig. S16). Interestingly, in the soil bacterium *Achromobacter,* the linker is also of a similar length and angle; the flagellin of *Sinorhizobium meliloti,* another soil dweller, also shows this linker but the angle is different. Our study suggests that in these species the linker may contribute to optimal adaptation of motility to different environments. Further structure-function studies may show by which mechanism rigid linkers in general contribute to motility across species.

Since both the linker and D4 domains individually emerged as affecting optimal motility, it was unexpected that deletion of the entire HVR did not compromise motility in soft agar assays to the same extent. This may be explained by the fact that functional flagella can be formed by flagellins that entirely lack D2, D3 or D4 domains, i.e., HVRs (e.g. *Bacillus subtilis[25]*), or harbor shorter HVRs: For example, *Pseudomonas aeruginosa* only contains a D3 and it adopts a highly elongated structure that also engages in interdomain contacts [26]. Thus, the significance of any or small (D3 only) vs. extended (D3 and D4) HVRs remains somewhat enigmatic and our data begs the question why EcN retains the biosynthetic burden of an additional ∼320 amino acids for an entire HVR that appears redundant for motility. We speculate that in the intestine, a greater adaptability to different environments – ranging from liquid environments to viscous intestinal mucus[27] – the HVR with a wider and much more textured outer surface provides the EcN flagellum with better adaptability. Future *in vivo* studies will be required to investigate this in detail.

In line with the TLR5 epitope mapping primarily to the D1 domain [13, 21], presence of linker, D4 or the HVR did not affect TLR5 recognition, at least *in vitro*. This indicates that the TLR5-mediated probiotic effect [6] is probably not dictated by unique HVR features of EcN flagellin directly, i.e. by modulating interaction with TLR5. Rather, we hypothesize that the effect may be indirect, e.g., EcN flagellins and resulting flagella may facilitate bacterial encroachment onto the epithelium by enhancing motility through the intestinal mucus. This may trigger more TLR5-signaling, which is required to mediate the probiotic effect. In line with this hypothesis, flagellin-dependent TLR5 signaling was shown to mediate the symbiotic properties of another gut commensal, *Roseburia hominis* [28] and thus, this phenomenon may be more common than previously thought. More extensive *in vivo* analyses are thus warranted.

That EcN flagellin can activate TLR5 but structurally would not permit formation of a (TLR5-EcN FliC)_2_ complex modelled on the (zebrafish TLR5-Salmonella FliC)_2_ [8] appears puzzling. However, recent data from zebrafish and grass carp indicate that the structure of *Salmonella*-TLR5 homodimers most likely represents an inhibitory complex [12], as TLR5a/TLR5b heterodimers (not homodimers as in the crystal structure) are required to sense flagellin in fish [11]. Thus, the structure of an activating complex with human TLR5 may generally differ from the only known structure to date. Depending on the features of the engaging flagellin [8], it might accommodate not only small HVRs but also longer HVRs connected to the D1 by a rigid linker. Further structural studies of co-complexes may be required to unequivocally confirm this.

Collectively, our study sheds light on structural features of the flagellin molecule of a primary human probiotic *E. coli* strain, enabling insights into the trade-offs faced by such bacteria in terms of immune recognition versus. motility in different environments. Clearly, environment-dependent motility appears more structurally constrained than immune recognition. Unless bacteria are prepared to accept complete loss of motility, TLR5 recognition accommodates enough structural flexibility to avoid more subtle attempts if immune evasion by bacteria that seek to retain motility. On the other hand, certain strains, e.g. *Campylobacter jejunii* [21], seem to have reached TLR5 ignorance whilst retaining motility. We hypothesize that in complex microbiomes trade-offs between immune recognition versus. motility are probably finely tuned and highly species-specific, cautioning against oversimplification. Our analysis highlights that a concomitant determination of structure-, motility- and immune-related parameters may be necessary for additional individual flagellins and their host species to deduce more general principles.

## Materials and Methods

### General materials

All used chemicals and reagents were purchased from Sigma unless otherwise stated. Kits are listed in Supplementary table S1.

### Gibson assembly, including generation of flagellin mutants

For the digestion of plasmid backbone, 2 µg plasmid were digested for 1 h at 37 °C using two of the following enzymes, ApaI, BSu15I (ClaI) and/or HindIII fast Digest (ThermoFisher), before enzyme activity was deactivated for 5 min at 95 °C. Digested plasmid fragments were separated on 0.8 % agarose gels, fragments of the desired size cut and DNA was isolated using GeneJET Gel Extraction Kit according to manufacturer’s manual. Fragments were dissolved in a volume of binding buffer equal to the weight (100 µl per 100 µg) at 60 °C for 10 min while shaking. DNA was bound to the column by centrifugation and then washed using the supplied Wash Buffer. After a dry centrifugation to remove excess liquid, the DNA was eluted using 30 µL ddH_2_O. The generated backbone fragment was then used for Gibson assembly. Gibson assembly was performed as published [29]. In brief, 25 µM primers (Supplementary table S2) with an overlapping region of 30 nucleotides to the target insertion site or another DNA fragment and 100 ng template DNA were used to amplify inserts using the SuperFi II Polymerase according to manufacturer’s instructions. The PCR run consisted of an initial denaturation at 98 °C for 30 s, followed by 35 cycles of 10 s at 98 °C for denaturation, 30 s at 60 °C for primer annealing and 30 s/kb at 72 °C for elongation before the final elongation for 10 min at 72 °C. The PCR products were applied to a 1% agarose gel and isolated from the gel as described for the digestion of the plasmid backbone. For the actual Gibson Assembly reaction, all fragments (2 – 3 µL of each) and the vector backbone (3 – 4 µL) were combined with 10 µL of Gibson Master Mix and incubated for 1 h at 60 °C. The generated target vector was transformed into a suitable bacterial host cell line (either *E. coli* DH5α or 116 λpir^+^) by heat shock. For this purpose, 6 – 10 µL of Gibson mix were incubated with 50 µL chemically competent bacteria for 30 min on ice, followed by heat shock at 42 °C for 40 s, 10 min on ice and incubation with 1 mL SOC medium at 37 °C for 1 h. Bacteria were plated on LB agar containing the suitable antibiotic. Colony formation was seen after an overnight incubation at 37 °C.

### Plasmid preparation

**For protein production and purification, EcN FliC constructs in the pET-19b (Novagen) vector were used. FliC_E, FliC_ETEV, FliC_**Δ**D0D1, FliC_**Δ**D4, FliC_**Δ**D4s and FliC_**Δ**D3D4 insert sequences. are listed in** Supplementary information**Supplementary figure legends** (included in the actual supplementary figures)

Amino acid sequences for constructs used. The constructs FliC_ΔD0 and FliC_ΔD0D3D4 were synthesized by Biocat (Heidelberg, Germany) and inserted in pET-19b vectors using restriction cloning with NcoI and BamHI. To reconstitute expression of FliCin fliC-deficient bacteria, a pGEM-T backbone was used. For modifying the chromosomal FliC region, pSB890 backbones were used. Plasmids used are standard subcloning, chemically competent E. coli DH5α cells were used (Invitrogen). Plasmids were isolated using commercial miniprep kits (Wizard Plus SV miniprep kit, Promega) and verified by automated DNA sequencing (Microsynth Seqlab).

### RNA isolation and quantitative PCR

RNA was isolated using the RNeasy Mini QIAcube protocol for animal tissue and cells with DNAse digestion according to manufacturer’s instructions. RNA was either directly reverse-transcribed into cDNA or intermediately stored at –80 °C. For reverse transcription,the High-Capacity RNA-to-cDNA-Kit (Thermo Fisher Scientific) was used according to manufacturer’s instructions, RNA concentrations were adjusted to 0.1 µg/µL. 10 µL of RT buffer and 1 µL of RT enzyme were added per sample and incubated for 1 h at 37 °C before the enzyme was inactivated for 5 min at 95 °C. 5 µL SYBR Green MasterMix (ROX, Roche), 0.3 µL of each primer (10 µM, Supplementary Table S2) and 1 µL undiluted (for hBD1 and hBD2 qRT-PCR) or 1:10 diluted (for GAPDH as housekeeping gene) cDNA were mixed with 3.4 µL ddH_2_O per reaction and run at a QuantStudio 7 Flex system. Thereby, an initial incubation at 50 °C for 2 min was followed by a denaturation step at 95 °C for 10 min. In the PCR stage, 40 cycles of a denaturation at 95 °C for 15 s alternated with an elongation at 60 °C for 1 min. To generate a melting curve, the samples were first heated to 95 °C for 15 s, then cooled to 60 °C, stepwise heated with 0.05 °C/s increments to 95 °C and kept at 95 °C for 15 s. Data was analyzed using the ΔΔC_T_ method.

### Bacterial culture

Bacteria were cultured in LB medium with selective antibiotics if not stated otherwise. Bacterial overnight cultures were set up from glycerol stocks, and day cultures by adding 50 µL overnight culture to 5 mL LB and grown while shaking at 120 rpm for 2 or 5 h for stimulation or motility assay, respectively. For quantification, bacteria were centrifuged at 3200 × g for 5 min, supernatants discarded, and bacteria resuspended in 1 mL PBS. Quantification was done by measuring the optical density (OD) at 600 nm assuming an OD_600_ of 1 contains 5 x 10^8^ colony forming units (CFUs) of *E. coli*. The concentration of bacteria was calculated according to the desired final multiplicity of infection (MOI).

### Soft agar motility assay

Soft agar plates were prepared from 5 g peptone, 2.5 g NaCl, 1.5 g agarose in 500 mL dH_2_O with 20 mL per Petri dish. Bacteria were cultured for 5 h in a day culture to reach a late exponential growth phase. 3 µl of day culture were poked into the center of a soft agar plate and incubated for 24 h at 37 °C. Pictures of the plates with a ruler were taken for quantification. The swimming radii were measured at three positions using ImageJ and averaged. To compare two strains on the same plate a pipet tip was dipped into the day culture and poked into the agar at approximately 1/4^th^ of the plate diameter. The same procedure was repeated with the second strain on the other half of the plate. Plates were incubated 24 h before pictures were taken. Diameters were not quantified due to strains hindering or influencing each other growth.

### Bacterial single cell swimming assay in liquid media

The *E. coli Nissle* strains WT, *fliC* Δ*D4*, *fliC* Δ*Linker*, *fliC* Δ*HVR* and Δ*fliC::FCF*, all containing a translational insertion in *malEK::PJ23119-mCherry* as a fluorescent marker were grown overnight in LB at 37 °C and 180 rpm agitation. The next day, strains were diluted to an OD_600_ of 0.15 and grown for 2 h at 37 °C and 180 rpm. OD_600_ was measured and samples were diluted 1:1000 in PBS supplemented with 0.2% of glucose (w/v) and carefully loaded in a self-made flow cell. Cell trajectories were recorded using a Ti-2 Nikon inverted microscope equipped with a 20× objective, by imaging the phase contrast channel at 20 fps for 10 s. Multiple videos were recorded and analyzed as follows: cell segmentation was performed in Fiji [30] using the MicrobeJ plugin, version 5.13p [31]. Subsequent analyses were done using custom Python scripts. For each detected particle, the centroid coordinates and area were extracted using the skimage.measure module from scikit-image [32]. Particles were filtered by area (5-150 pixels) to remove false detections. Tracking was performed on the resulting binary masks using the Python package trackpy, version 0.6.4, with a search range of 5 pixels and a memory of 1 frame. For each tracked particle, centroid positions were used to calculate instantaneous velocities from frame-to-frame displacements divided by the time interval (50 ms). The mean velocity for each track was calculated as the average of its instantaneous velocities. Only tracks with durations of at least 3 s were included in the analysis. Based on the non-motile Δ*fliC* control, tracks with mean velocities below 5 µm/s were considered non-motile (Brownian motion) and excluded from velocity statistics. For each strain, data were pooled from multiple regions of the flow cell.

### Negative stain transmission electron microscopy (TEM) of bacteria

For negative stain TEM of flagellated bacteria, bacterial day culture was inoculated with 500 µL overnight culture in 10 mL LB media. Bacteria were grown while shaking at 120 rpm, 37 °C for 4 h, adjusted to 1 × 10^9^ CFU/mL and transferred to the Electron Microscopy Facility at the Max Planck Institute for Biology Tübingen. The bacteria were shaken for another 1 h before 1 – 1.5 mL of culture were spun down at 5000 × g for 3 min and the pellet was resuspended in 100 µL of culture. 4 µL of resuspended bacteria were applied to glow-discharged carbon-coated EM grids and incubated for 2 min. The grid was washed with three drops of ddH_2_O and three drops of 1% uranyl acetate. The 4^th^ drop of 1% uranyl acetate was incubated for 5 min before the liquid was removed and the grid was dried. Grids were analyzed in a transmission electron microscope (Tecnai Spirit, Thermo Fisher Scientific, Eindhoven, The Netherlands) equipped with a CMOS camera (TemCam-XF416, TVIPS, Gilching, Germany).

### Negative stain transmission electron microscopy (TEM) of isolated flagella

For negative-stain electron microscopy of isolated flagella, overnight cultures were used to inoculate 100 ml LB medium to an OD_600_ of 0.01. Bacteria were grown at 37 °C with shaking at 80 rpm overnight. Cells were harvested by centrifugation at 2,000 × g for 20 min at 4 °C, and the supernatant was discarded. The pellets were resuspended in 10 ml of HEPES (20 mM, 150 mM NaCl, pH 7.4) buffer per sample. To shear off the flagella, the bacterial suspensions were transferred to new flasks and stirred at 500 rpm for 1 h at 4 °C. Bacterial cells were removed by centrifugation at 8,000 × g for 30 min at 4 °C, repeated twice. The supernatants were collected and concentrated using ultrafiltration units (Amicon 10 kDa MWCO). Centrifugation was performed at 3,214 × g at 4 °C. The concentrated flagella suspension was stored at 4 °C. For negative staining, 4 µl of the isolated flagella suspension was applied to glow-discharged carbon-coated grids for 1 min and gently blotted away with filter paper. This procedure was repeated three times with ddH_2_O. Subsequently, 4 µl 2% (w/v) phosphor-tungstic acid was applied for 45 s, replaced with 4 µl ddH_2_O, and liquid was directly removed by blotting with filter paper. The grids were allowed to air dry for 5 min. Samples were examined using a JEOL JEM-2100 Plus transmission electron microscope operated at 200 keV. Images were acquired at various magnifications. Flagella diameters were measured using (Fiji is just) ImageJ (v1.54). Prior to measurements, background subtraction was performed using the rolling ball method (2 px radius), and diameters were determined from plot profiles.

### Silver staining of SDS-PAGE gels

For visualization of low protein concentrations, the Pierce Silver Stain Kit (Thermo Fisher) was used according to the manufacturer’s instructions.

### Protein production and purification for structural studies

Pre-cultures were inoculated with *E. coli* BL21 containing the appropriate plasmid and grown overnight. 1-6 L LB media was inoculated 1:100 with overnight culture and grown at 37°C while shaking until an OD_600_ of 0.6. Protein production was then induced with 0.2-0.5 mM IPTG, and cells were grown at 25 °C overnight or 37 °C for 4 h. Cells were harvested by centrifugation for 15 min at 9200 × g (Sorval RC 6+, Thermo Fisher Scientific). The pellets were frozen in liquid nitrogen and stored at -80 °C. Cell pellets were resuspended in 5 mL lysis buffer (50 mM Tris pH 8.0, 300 mM NaCl) per gram pellet wet-weight. The suspension was supplemented with 1x “cOmplete” protease inhibitor (Roche, Basel, Switzerland) and 125-500 U Benzonase Nuclease (Merck KGaA, Darmstadt, Germany). Cells were lysed by sonication with an amplitude of 40%, for 3 min, with on-off pulse cycles of 0.5 s (Digital Sonifier 250, Branson Ultrasonics). The lysate was centrifuged for 45 min at 34’500 × g (Sorval RC 6+, Thermo Fisher Scientific,), filtered (0.22 µm cutoff) and loaded on a 5 mL HisTrap FF crude column (GE Healthcare) equilibrated with His-A buffer (50 mM Tris pH 8.0 300 mM NaCl) using an ÄKTAprime plus FPLC system (GE Healthcare). After sample loading the column was washed with His-A and 2% His-B Buffer (50 mM Tris pH 8.0, 300 mM NaCl, 500 mM imidazole). Proteins were eluted using 10%, 20%, 30%, 60% and 100% His-B Buffer steps. Fractions were pooled based on SDS-PAGE analysis. Pooled protein fractions were dialyzed against SEC-buffer, or His_A_ buffer, using a Spectra/Por dialysis membrane at 4° C overnight (Spectrum Laboratories Inc). If proteolytic cleavage of the His-tag was attempted, 1-2 mL TEV protease (0.5 mg/mL, produced in our laboratory) was added to the pooled fractions, before dialysis. The His-tagged TEV protease and undigested protein were removed by a second IMAC. The flow through was pooled and yield and purity assessed by SDS-PAGE. For size exclusion chromatography, pooled samples were filtered (0.22 µm) and loaded on an equilibrated SEC-column. For preparative SEC a HiLoad 16/60 Superdex 200 or Superdex 75 column (Pharmacia) with a Pharmacia LKB GP250 plus system were used. For analytical SEC a 3.2/30 Superdex 200 Increased column (GE Healthcare) or a 3.2/30 Superdex 75 column (Pharmacia) was used with an Ettan LC system (GE Healthcare). For crystallization a SEC-buffer containing 20 mM HEPES pH 7.5 and150 mM NaCl was used. For stability experiments DPBS-buffer (0.90 mM CaCl_2_, 0.49 mM MgCl_2_, 2.67 mM KCl, 1.47 mM KH_2_PO_4_, 137.93 mM NaCl, 8.06 mM Na_2_HPO_4_) was used for SEC.

### Crystallization

To determine an appropriate starting concentration for initial crystallization experiments, a precipitation test was done. 0.5 µL of filtered (0.22 µm) protein solution was mixed on a glass slide, with different precipitate (30% (w/v) PEG4000 and 3 M ammonium sulfate) solutions. The protein concentration was stepwise increased until the protein precipitated within 1 min with at least one of the two precipitants. For initial screens, commercially available kits were used according to the manufacturers’ instructions: JCSG, Morpheus (Molecular Dimensions), Wizard I-IV (Emeraly BioSystems), Crystal Screens I and II and PEG ION (Hampton Research). Crystallization screens were done using 96-well siting drop Intelli-plate (Art Robbins Instruments, Sunnyvale, USA) with either a Gryphon (Art Robbins Instruments, Sunnyvale, USA), or Freedom EVO (Tecan, Männedorf, Switzerland) crystallization robot. Drops were set up in a one-to-one ratio of protein solution and mother liquor, with a total drop size of 400 nL with the Gryphon, and 600 nL with the Freedom EVO. Crystals were optimized using 4 × 6 (Hampton Research, Aliso Viejo, USA) hanging drop plates.

### Structure determination

To optimize the cryo-protectant or to test diffraction capacity of a crystal, the in-house X-ray system was used. This system is composed of a rotating copper anode X-ray source (MicroMax-007HF (Rigaku, Sevenoaks, UK) producing CuKα radiation at λ = 1.5418 Å and a mar345 image plate detector (marresearch, Norderstedt, Germany). The HATODAS II [33] server was used to identify promising heavy atoms for initial soaking experiments. Therefore, crystals were soaked for 5 and 10 min in crystallization solutions containing different heavy atoms with a concentration of 10 mM or saturated if the solubility was lower. For Sm^3+^ and UO_2_^2+^ compounds various crystals were soaked with concentrations ranging from 2.5 mM to 10 mM and soaking time between 1 min and 24 h. Additionally, crystals were soaked with 3 mM (NH_4_)_2_WS_4_ or Lu(Ac)_3_ for 2 h, 1 mM Ta_6_Br_14_ for 1 week, 2.5 or 5 mM EuCl_3_ for 24 h or 1 mM SeC(NH_2_)_2_ for 10 min. Data sets were collected at the macromolecular crystallography beamline X06DA-PXIII of the Swiss Light Source (Paul Scherrer Institute, Villigen, Switzerland). Data sets for structure determination via molecular replacement were collected with λ = 1 Å (12.3984 keV). For anomalous data the wavelength was adjusted to specific values, according to the absorption edge of the element used. The XDS program package was used to process (index, integrate, scale) the experimental diffraction image data sets, XSCALE to scale and merge datasets, XDSCONV to convert different crystallographic file formats [34]. From the PHENIX [35] program package XTRIAGE [36] was used to analyse data sets and PHENIX.REFINE [37] was used for simulated annealing and refinement. From the CCP4 [38] program package, POINTLESS [39] was used to identify possible space groups, MATTHEWS_COEF [40] was used to evaluate the possible unit cell content, CHAINSAW [41] to modify models for molecular replacement, PHASER [42] and MOLREP [43] for molecular replacement, REFMAC5 [44] for refinement and COOT [45] for model building and evaluation. PYMOL [46] was used for structure examination and figure generation. Experimental phasing was done using SHELX C/D/E [47] and AUTOSHARP [48] with SOLOMON [49] for density modification. MOLPROBITY [50] was used for model evaluation. The following structure files were deposited to rcsb.org: [to follow]

### Cultivation and stimulation of mammalian cells

HEK293T cells were grown in DMEM supplemented with 10% FCS and 1% Penicillin/Streptomycin. mTLR5 HEK293T cells required the selective antibiotics Normocin (50 mg/L) and Blasticidin (10 mg/L), while hTLR5 HEK293T cells required Blasticidin (10 mg/L), Hygromycin B (100 mg/L) and Tetracycline (12.5 mg/L) that were added freshly to the medium. CaCo-2 (subclone C2BBe1) cells were cultured in DMEM supplemented with 10% FCS, 1% non-essential amino acids (Gibco) and 1% Penicillin/Streptomycin. THP-1 cells were cultured in RPMI 1640 (very low endotoxins), supplemented with 10% FCS, 1% L-glutamine (gibco) and 1% Penicillin/Streptomycin. Murine bone marrow derived dendritic cells (BMDCs) were cultured and differentiated in RPMI 1640 (very low endotoxins), supplemented with 10% FCS, 4% GM-CSF conditioned medium (produced in our laboratory), 1% non-essential amino acids (gibco), 1% Penicillin/Streptomycin and 0.5% β-mercaptoethanol. Cells were incubated at 37°C 5% CO_2_ and split every 3 - 4 days. **Error! Reference source not found.**2 × 10^5^ cells per well in a 24-well plate or 3 × 10^4^ cells per well of a 96 well plate were seeded in the respective culture medium. For stimulation with recombinant FliC, cells were incubated ∼ 40 h at 37 °C 5% CO_2_ before medium was replaced by medium containing recombinantly isolated FliC proteins for 24 h. In case of bacterial infection, medium was replaced 24 h after seeding with antibiotics-free medium and cells were incubated overnight at 37 °C 5% CO_2_ before medium was exchanged to antibiotics-free medium containing bacteria for a total of 24 h or 6 h in case of C2BBe1 cells. 1 h post infection (p.I.) gentamicin was added to inhibit overgrowth of bacteria. For both stimulation and infection, supernatants were taken, centrifuged at 3200 × g for 10 min, or max speed for 6 min and transferred to a new tube or plate. Cell-free supernatants were used for ELISA analysis and C2BBe1 cells were lysed in 350 µL of RLT/1% β-mercaptoethanol and frozen at –80 °C for RNA isolation. To differentiate THP-1 cells to a macrophage-like phenotype, 2 × 10^5^ cells per well of a 24-well plate were seeded in medium containing 100 nM PMA. Three days later, PMA was removed and cells were left to rest for 72 h at 37 °C 5% CO_2_ before stimulation/infection (as described above).

### Cultivation and stimulation of enteroids

To revive enteroids, cells were washed twice with pre-warmed DMEM before being in ice-cold Matrigel. 4 drops of 10 µL Matrigel-enteroid mix were placed into one well of a 24 well plate. A 10 min incubation at 37 °C followed to polymerize the Matrigel. 400 µL of enteroid passage medium (growth medium [Advanced Dulbecco’s modified Eagle medium/F12, 1 mM HEPES, 1x Glutamax, 1x B27, 1 mM N-Acetylcysteine, 10 nM Gastrin, 50 ng/mL EGF, 10 mM Nicotinamide, 500 nM A83-01, 10 μM SB202190, 50% L-WRN-conditioned media (contains Wnt3a, R-spondin 3, and Noggin)] + 10 μM Y27632, 250 nM CHIR99021) were added. 2 – 3 days afterwards the medium was replaced by enteroid growth medium.

To split enteroids, Matrigel was dissociated by 1 mL of TrypLE express (gibco) and careful pipetting ensured breaking. The cell-Matrigel mix was diluted 1:10 and incubated at 37°C while pipetting until single cell dissociation was achieved. To inhibit trypsin, 100 µL of FCS were added. Enteroids were pelleted at 300 × g for 5 min and resuspended in Matrigel and dropped into a 24 well plate and cultivation was performed as described above. The medium was replaced every 2 - 3 days by fresh enteroid growth medium and cells were split every 1 – 2 weeks, when they grew large or dense enough. For stimulation, the enteroids were treated as if they were split. When enteroids in fresh Matrigel were distributed in 96 well plates, one 10 µL drop was placed in one well and 200 µL of enteroid passaging medium was added. 2 - 3 days later it was replaced by 200 µL of enteroid growth medium. Enteroids were in culture for at least 4 days prior to stimulation. For stimulation, the medium was replaced by recombinant FliC containing medium and left on cells for 18 h. Supernatants were taken and used for IL-8 ELISA.

### ELISA

Generally, ELISAs were performed according to manufacturer’s protocols using commercially available kits (Supplementary Table S1). If necessary, buffers were prepared accordingly. Dilutions of reagents used were done as recommended by the manufacturer in suitable buffers. Generally, sandwich ELISAs were performed. Thus, the plates were coated with a suitable capture antibody overnight at 4 °C, before the coating solution was washed off in four consecutive washing steps and replaced by the double standard volume (e.g. 200 µL or 80 µL) of the suitable blocking solution. All following incubations were at room temperature while swaying. The blocking buffer was removed, the plates washed four times and the standard protein, as well as the cell-free samples in a suitable dilution were applied and incubated. Another four times wash followed, and the detection antibody fused to biotin was incubated on the wells. After four more washes, the standard volume of a streptavidin-tagged alkaline phosphatase (AP) or horseradish peroxidase (HRP) was added to the wells. An incubation followed before the conjugate of enzyme and streptavidin was removed by a more vigorous washing step according to the manufacturer’s instructions. The substrate, either para-nitrophenyl phosphate (pNPP) for AP or tetramethylbenzidine (TMB) for HRP, was prepared and added to the wells. The incubation was performed in the dark at room temperature without swaying for 20-30 min until the desired color change was reached. The reaction was either stopped or not using 2N sulfuric acid. The absorption was read at a wavelength of 405 nm for AP and 450 nm for HRP using a spectrophotometer.

### Flow cytometry

For flow cytometric analysis, cells were carefully scraped off the wells and transferred to a V-bottom 96 well plate and unspecific binding was blocked by adding 100 µL of either cell culture supernatants of 2.4G2 cells secreting an antibody directed against the murine Fcγ receptor for murine cells or a commercially available human Fc-Block (BD; diluted 1:20 in FACS Buffer) for human cells for at least 15 min at room temperature. The cell pellet was washed twice using PBS to remove FCS before staining for their viability using a fixable viability dye (ThermoFisher) diluted 1:10,000 in PBS for another 15 min in the dark on ice or at 4°C. After two more washing steps in PBS 1% FCS the staining solution containing the desired antibodies in a 1:100 dilution (α-mCD11b BV421 (HL3, BD), α-mMHC-II PerCp-cy5.5 (M5/114.15.2, BD), α-mCD86 PE-cy7 (GL1, BD), α-mCD40 APC (3/23, BD), α-mTLR5 PE (ACT5, BD) was added to the cells and incubated for 30 min in the dark at 4 °C or on ice. The cells were washed another one time before resuspending them in 100 µL of FACS Buffer. Measurements were done on a BD LSR Fortessa II. Analysis was done using FlowJo v10.

### Animals

C57BL/6J and C57BL/6N mice (Charles River Laboratories, Sulzfeld, Germany) were ordered at an age of 6-8 weeks at least one week prior to start of the experiments. Mice were housed under specific-pathogen-free (SPF) conditions in individually ventilated cages (IVCs). All experiments were carried out in compliance with specific local institutional guidelines on animal experiments under regular hygiene monitoring in accordance with the German animal protection law and according to regulations by Gesellschaft für Versuchstierkunde/Society for Laboratory Animal Science, GV-SOLAS, as well as according to the European Health Law of the Federation of Laboratory Animal Science Associations, FELASA. All experiments were registered and approved by the local federal authority, the Regierungspräsidium Tübingen, under approval H01/20M.

### Isolation, differentiation and stimulation of primary bone marrow derived dendritic cells

Bone-marrow derived dendritic cells (BMDCs) were isolated and differentiated according to a published protocol [51]. Therefore, mice were sacrificed at an age of about 8 weeks, femurs and tibias were isolated, freed of residual tissue and sterilized using ethanol while still intact. Bones were cut open at both ends under sterile conditions, and a 0.45 mm syringe filled with sterile PBS was used to flush out the bone marrow cells. Cells were spun down at 300 × g for 5 min, supernatant discarded and washed using fresh PBS. 2 × 10^6^ cells were seeded per 10 cm Petri dish (not cell culture treated) in 10 mL of GM-CSF containing BMDC culture medium as listed above**Error! Reference source not found.**. The cells were incubated at 37°C 5% CO_2_ for two days before another 10 mL of fresh BMDC culture medium was added to feed the cells. Three days after the feeding, 10 mL of the culture supernatant was taken off the cells, centrifuged at 300 × g for 5 min, supernatant was discarded, and cells resuspended in 10 mL fresh BMDC culture medium. Cells were subsequently transferred back to the Petri dish and incubated two more days. To seed the cells for stimulation, cells were pooled and 2 × 10^6^ cells per well were seeded in a 6-well plate 1 h prior to stimulation in BMDC culture medium. Either recombinantly isolated proteins at the indicated concentration or bacteria grown for 2 h in a day culture were added at an MOI of 2. BMDC culture medium contains penicillin and streptomycin, thus no gentamicin kill was additionally performed. BMDCs were incubated for 16 h with FliCs or bacteria before supernatants were taken, centrifuged at max speed for 5 min and transferred to a new tube. Cells were scraped off at 16 h after incubation for flow cytometric analysis.

### Statistical analysis

Collected data was visualized and statistically analyzed using GraphPad Prism version 8 and 10. First, Shapiro-Wilk test was deployed to test for normality in each dataset (e.g,. each concentration of each construct or each MOI of each bacterium separately). If only few datasets (less than 20% of datasets) were not normally distributed according to Shapiro-Wilk test, normality was nevertheless assumed because of small sample sizes (often n=3), and when normal distribution in other data sets of same stimulus often only varying in concentration/MOI was confirmed. ANOVA, either ordinary one-way or 2-way ANOVA or Kruskal-Wallis test for not-normally distributed data with Dunnett correction for multiple testing, was chosen depending on the nature of the data to be analyzed as indicated in the figure legends. Considering p values, the following scheme was applied: * p ≤ 0.05, ** p ≤ 0.01; *** p ≤ 0.001; **** p ≤ 0.0001. In some cases, two statistical tests were performed on the same data set. One test was utilized to test for statistical differences of the sample compared to unstimulated control (mock). Here, the same significance levels as described above were used, but the symbol used was ‘#’ instead of ‘*’. The ‘*’ symbol was used to indicate statistical differences between a given sample and either EcN WT, in case of bacterial infection, or full-length EcN FliC, in case of stimulation with recombinantly isolated proteins. Again, the same significance levels as described above were used with an ‘*’ symbol. The test used, the group to which the sample was tested against (sample versus mock or sample versus EcN WT or sample versus full-length) as well as the symbol used (# or *) to depict significance level are indicated in the figure legends.

## Acknowledgements

We thank Libera Lo Presti for excellent editorial support.

## Author contributions

(according to Credit guidelines)

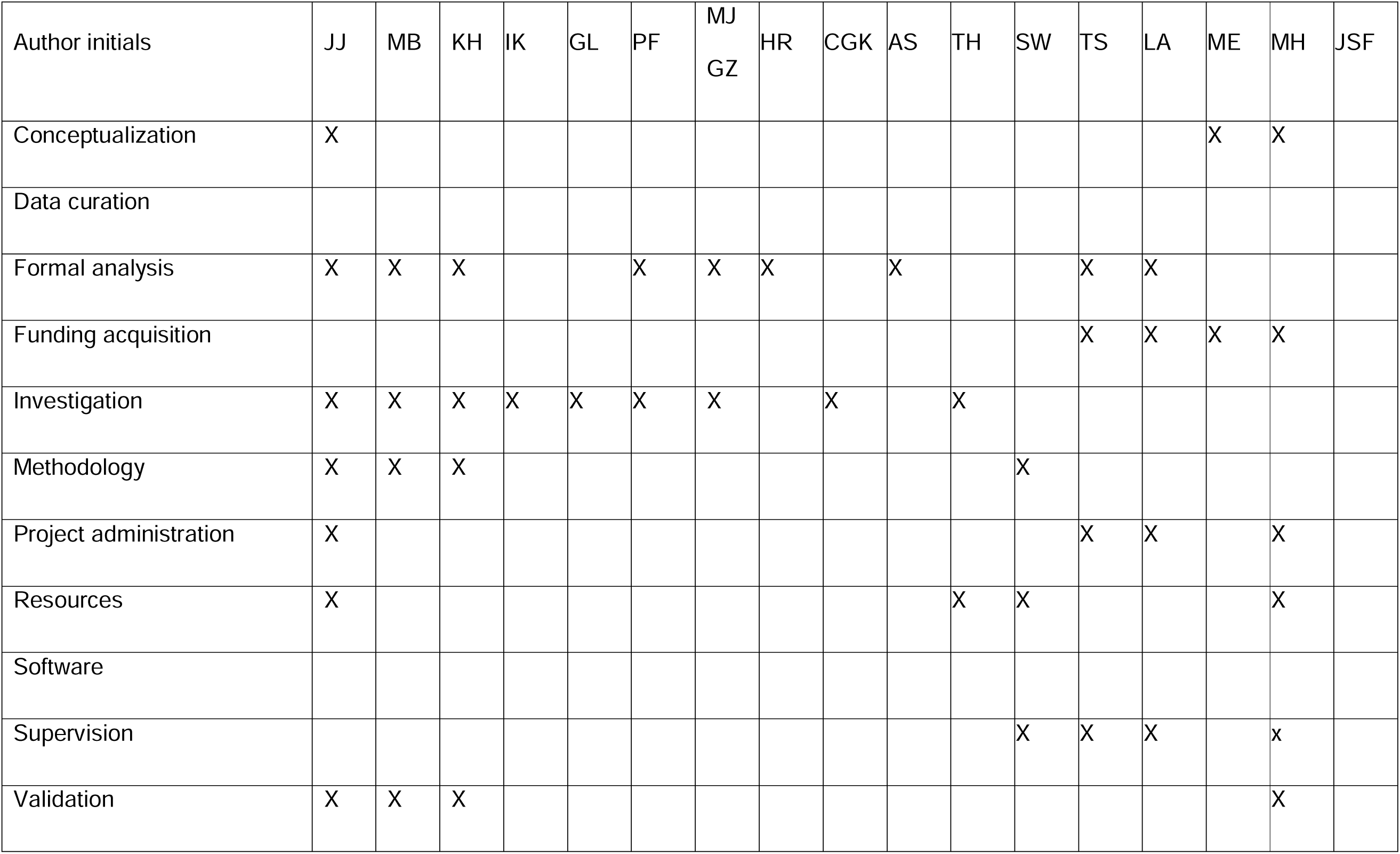
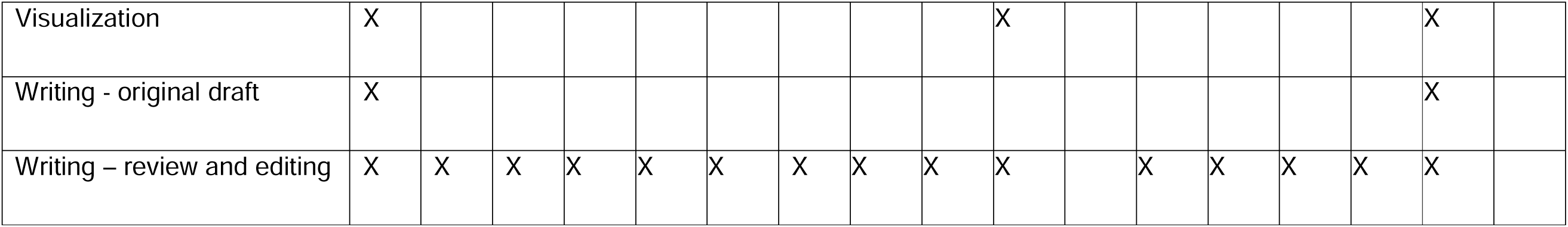

## Supplemental information

**Supplementary figure legends** (included in the actual supplementary figures)

**Amino acid sequences for constructs used**

All constructs for protein purification contained EcN FliC sequences fused N-terminally to an 8xHis tag (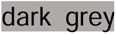) connected by a linker (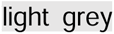) containing a TEV cleavage site (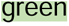). Domain assignment is shown exemplarily for the full-length EcN FliC. 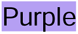 marks the N and C-terminal D0 domain, 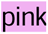 the D1 domain, 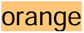 the linker region, 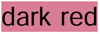 the D2 domain, 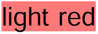 the D3 domain and 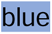 the D4 domain. In the ΔD4 construct, a short linker is inserted to connect the N- and C-terminal parts of the D3 (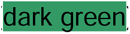).

Full-length EcN FliC:

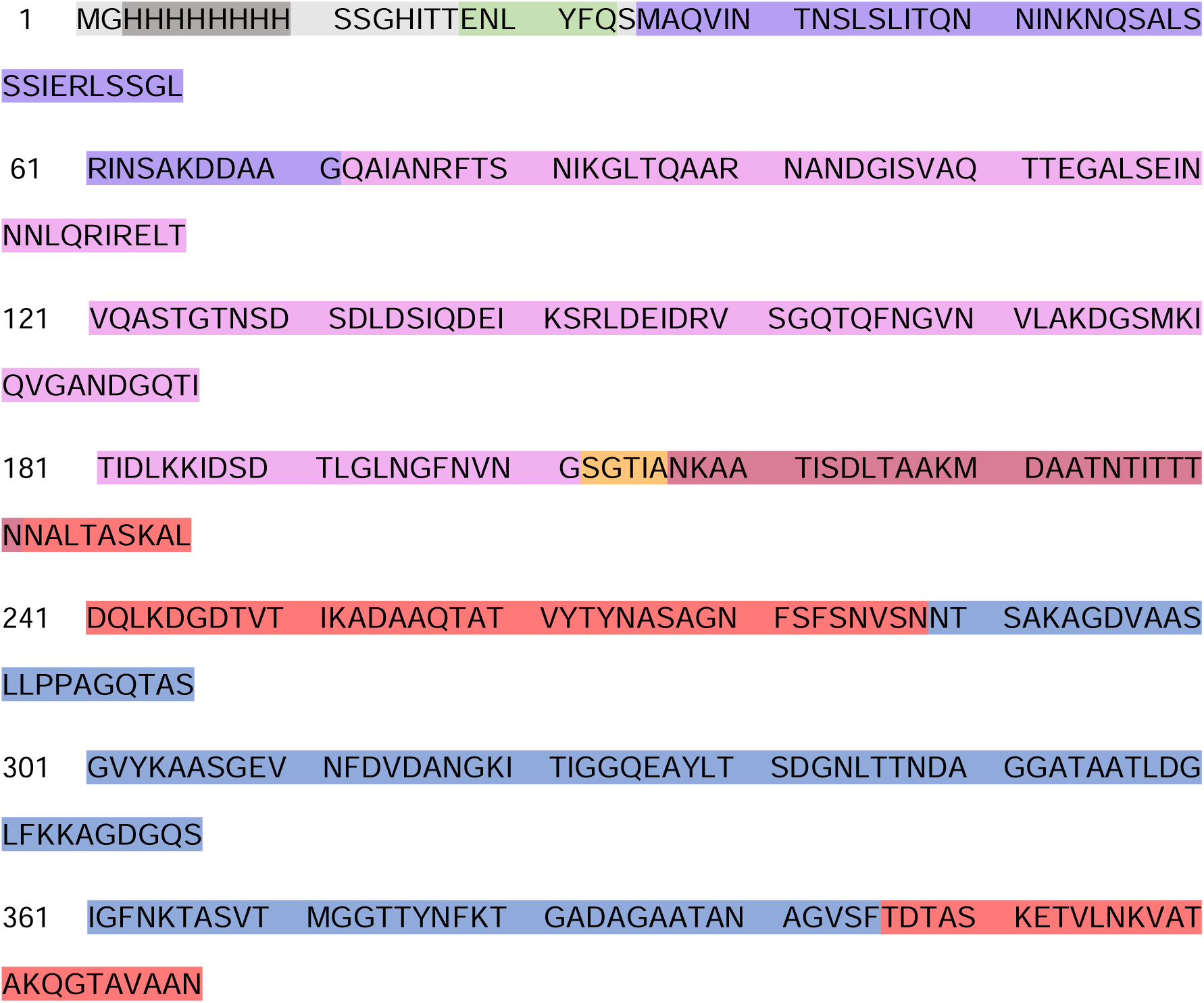
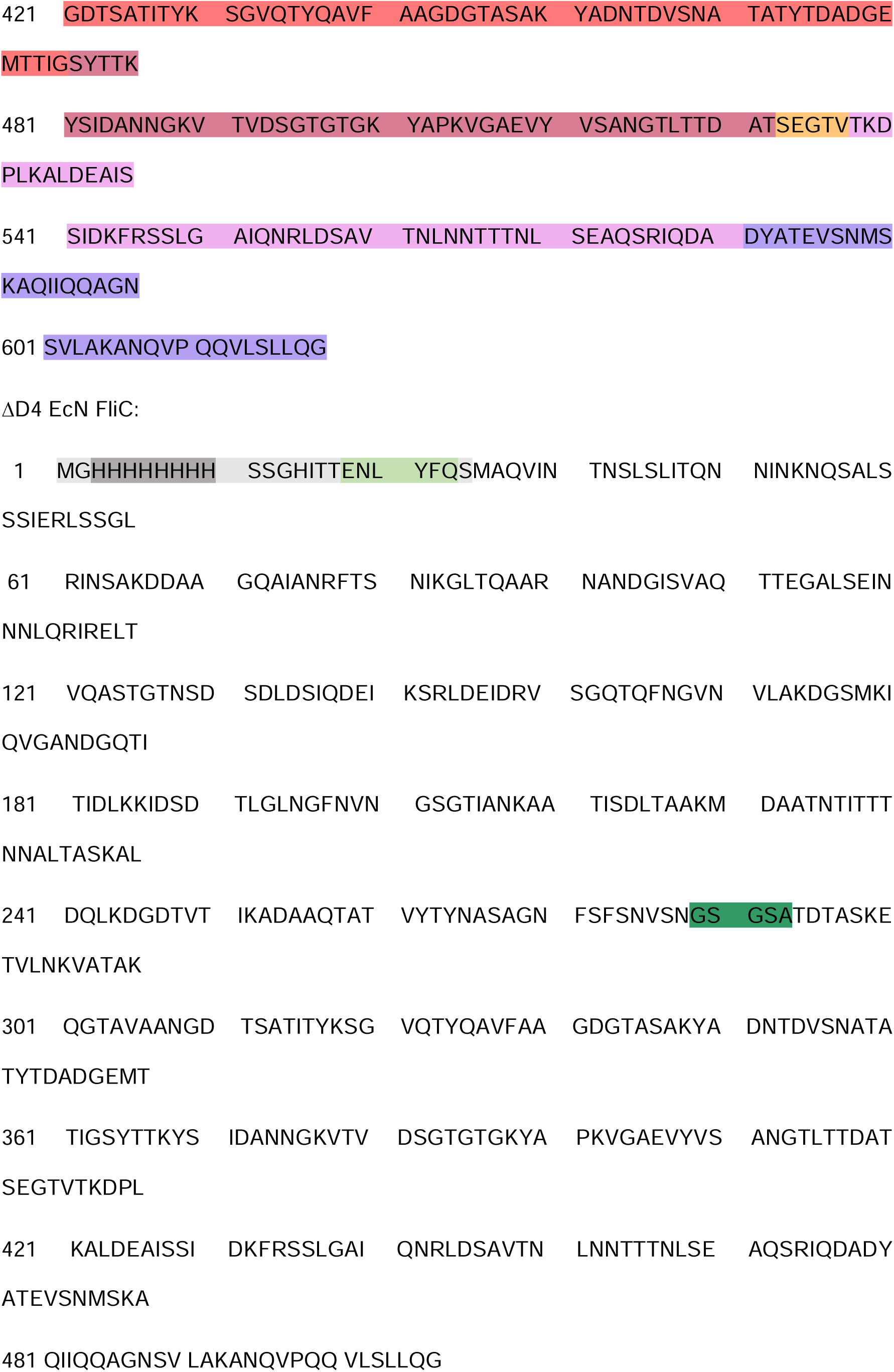
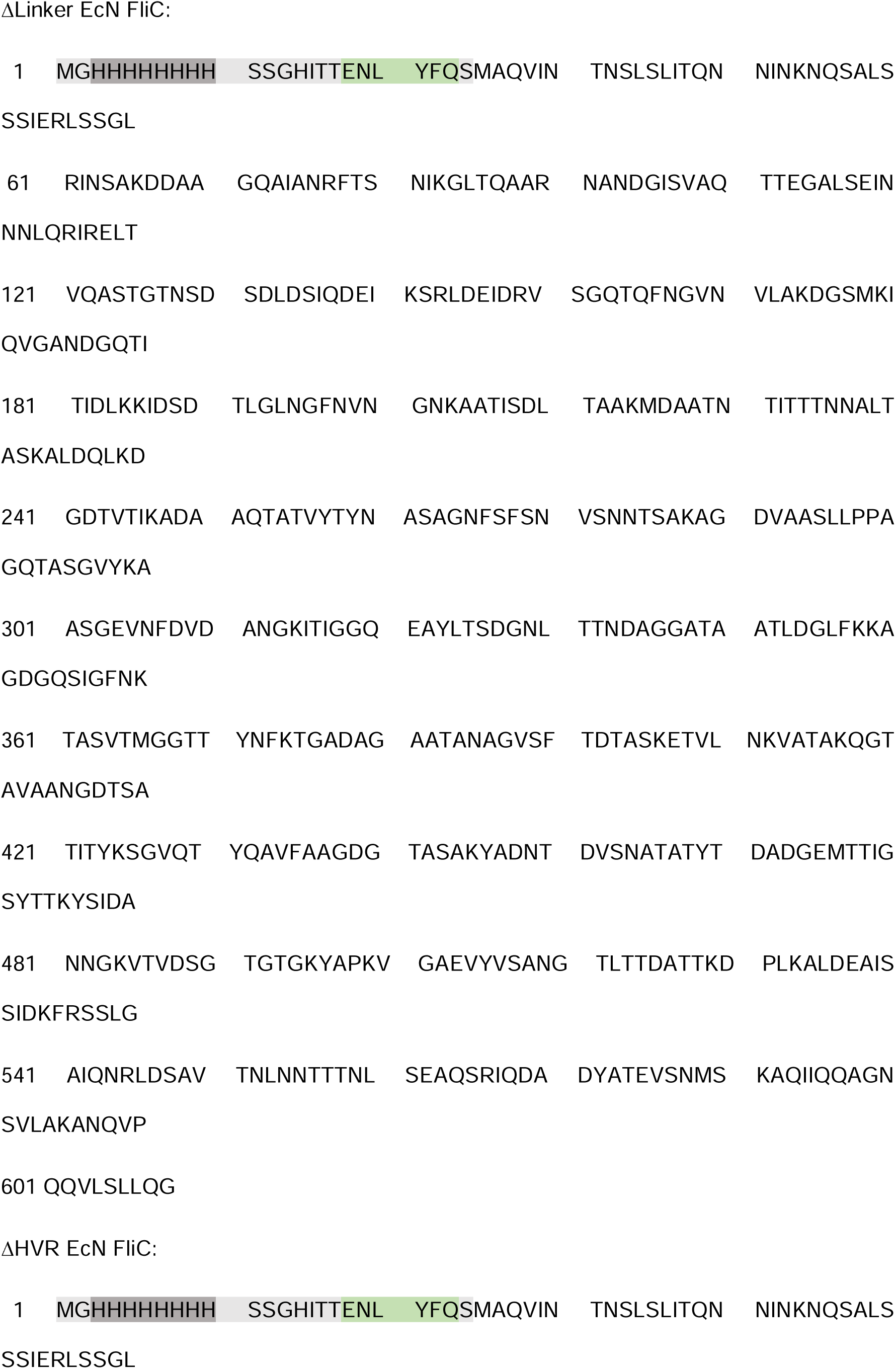
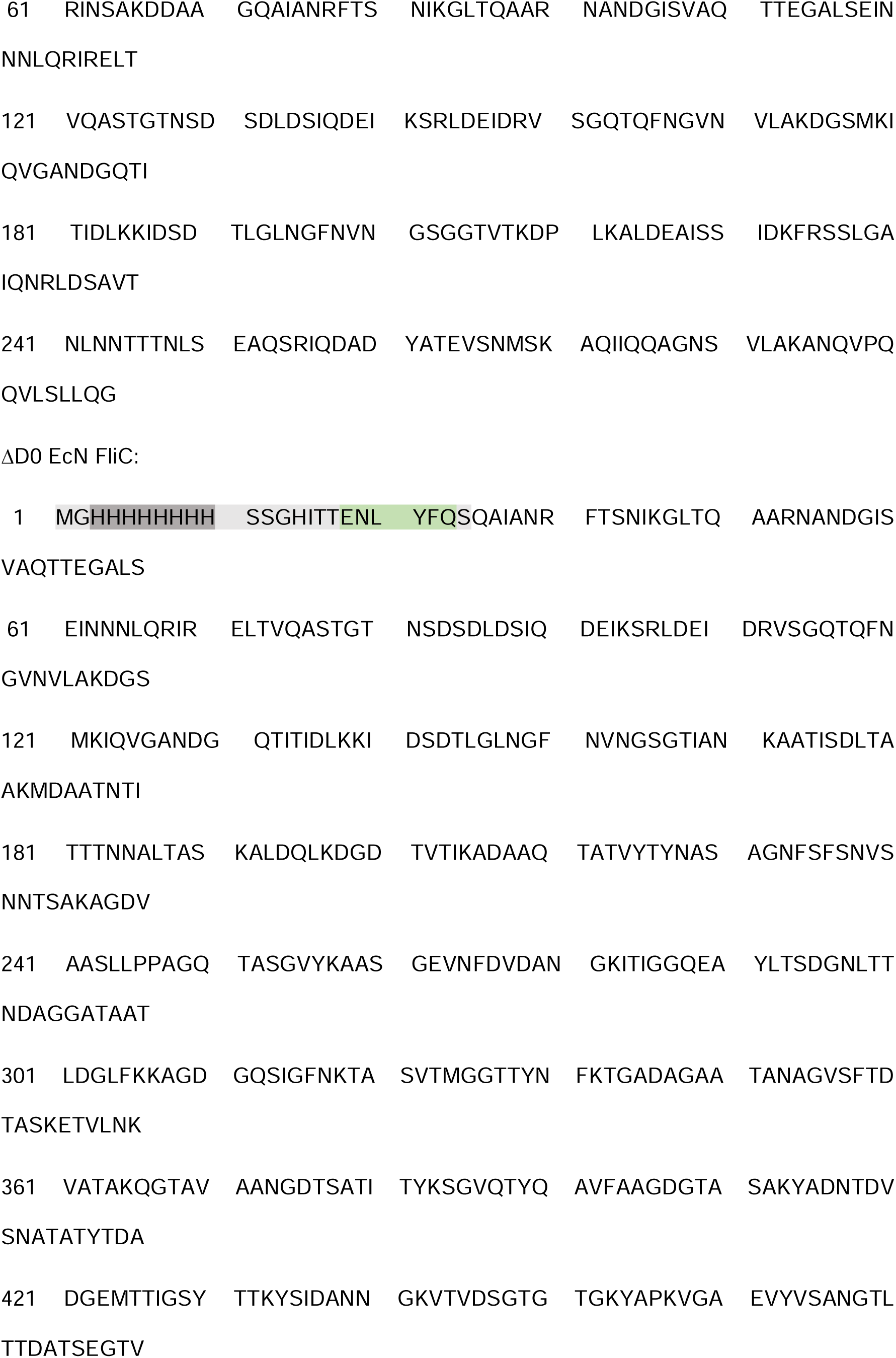
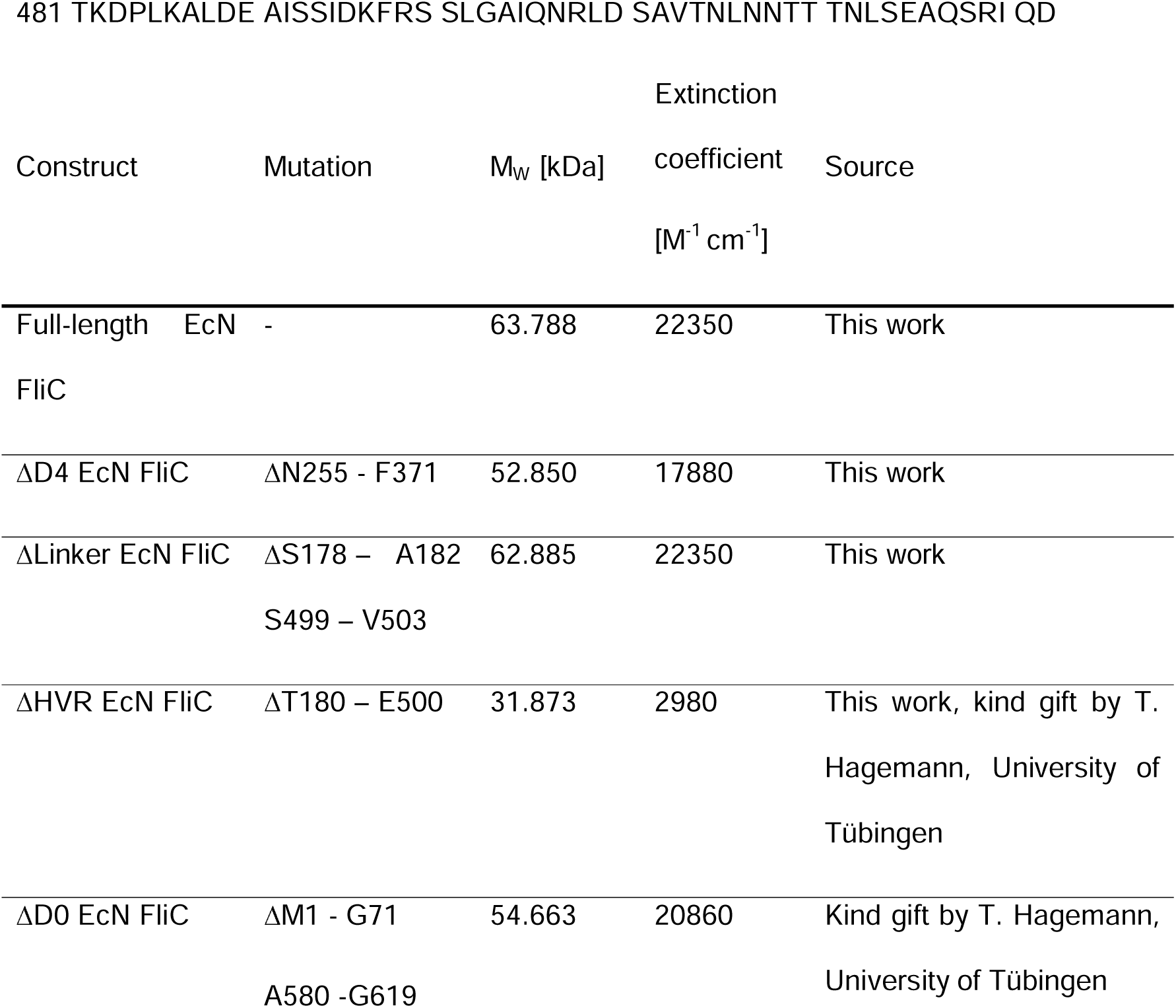

## Supplementary tables

**Supplementary Table S1:** Used kits.

| KIT | Manufacturer | Catalog # |
| --- | --- | --- |
| ELISA MAX™ Deluxe Set Human IL-8 | BioLegend | 431504 |
| GeneJET Gel Extraction KIT | Thermo Fisher Scientific | K0692 |
| GenElute bacterial genomic DNA isolation Kit | Merck | NA2100 |
| Gibson assembly MasterMix | NEB | E2611S |
| High-Capacity RNA-to-cDNA-Kit | Thermo Fisher Scientific | 4387406 |
| Human IFNβ ELISA | R&D | DY814-05 |
| Human IL-10 ELISA | BD | 555157 |
| Human IL-1β ELISA | R&D | DY201-05 |
| Human IL-6 ELISA | R&D | DY206 |
| Human TNF ELISA | BD | 555212 |
| Invitrogen Platinum SuperFi II Polymerase + 5 X Buffer | Thermo Fisher Scientific | 12361050 |
| Murine IL-10 ELISA | R&D | DY417-05 |
| Murine IL-1β ELISA | R&D | DY401-05 |
| Murine IL-6 ELISA | BD | 555240 |
| Murine TNF ELISA | BD | 555268 |
| Pierce™ Silver Stain Kit | Thermo Fisher Scientific | 24612 |
| PureYield Plasmid Mini Kit | Promega | A1222 |
| PureYield™ Plasmid Midiprep System | Promega | A2496 |
| RNAse-free DNase Set (250) | QIAGEN | 79256 |
| RNeasy Mini QIAcube Kit | QIAGEN | 74116 |

**Supplementary table S2:**
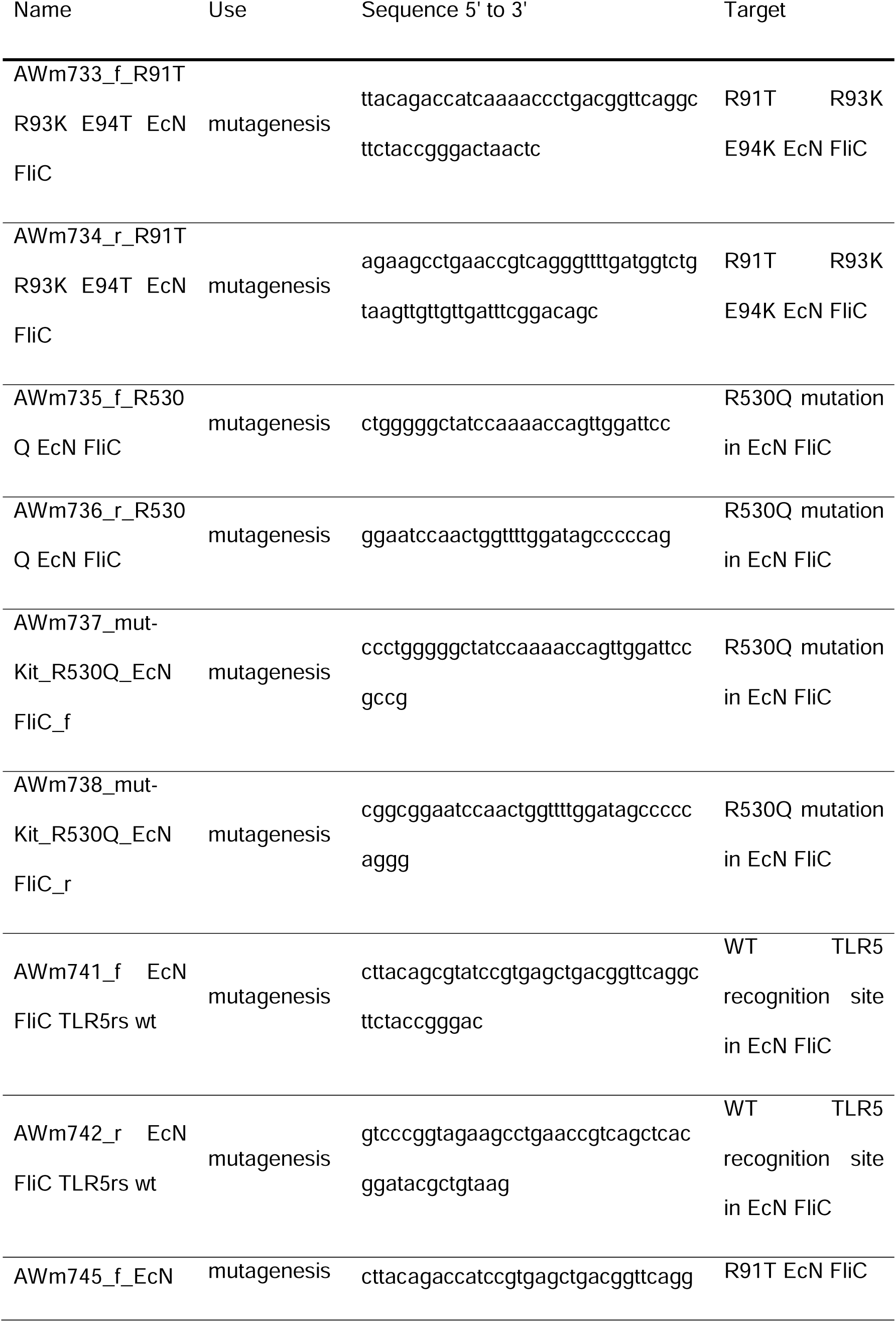
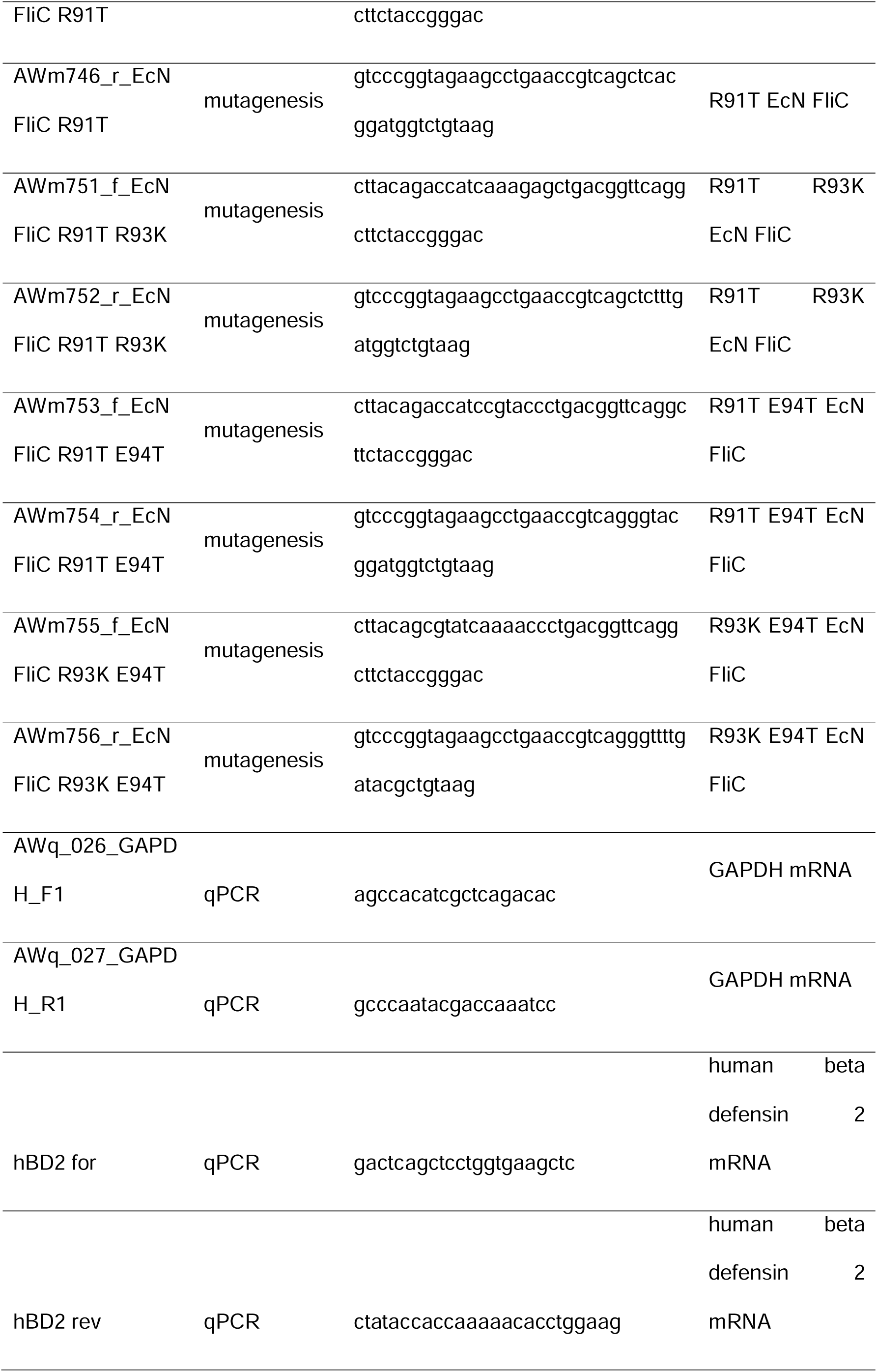
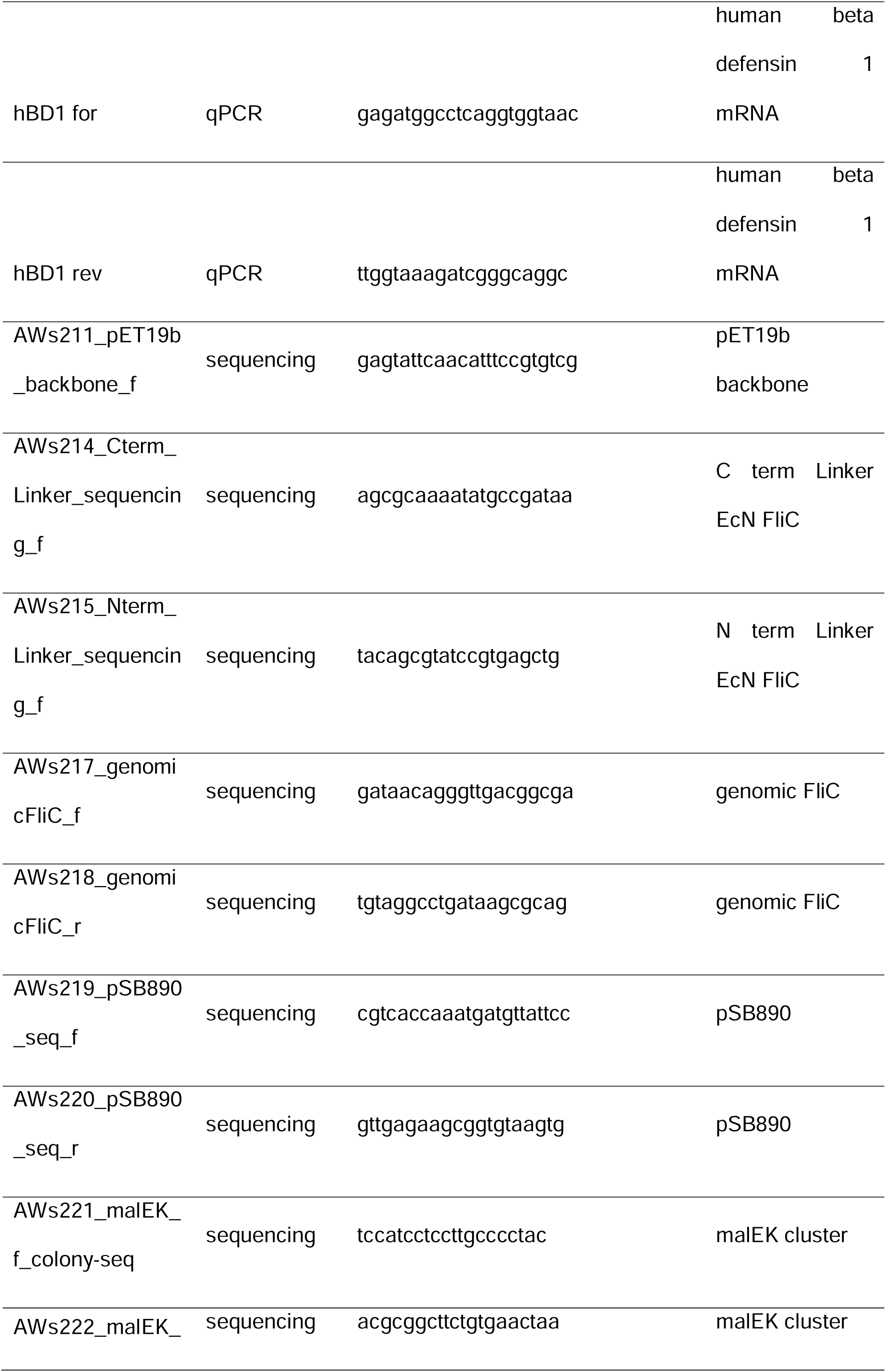
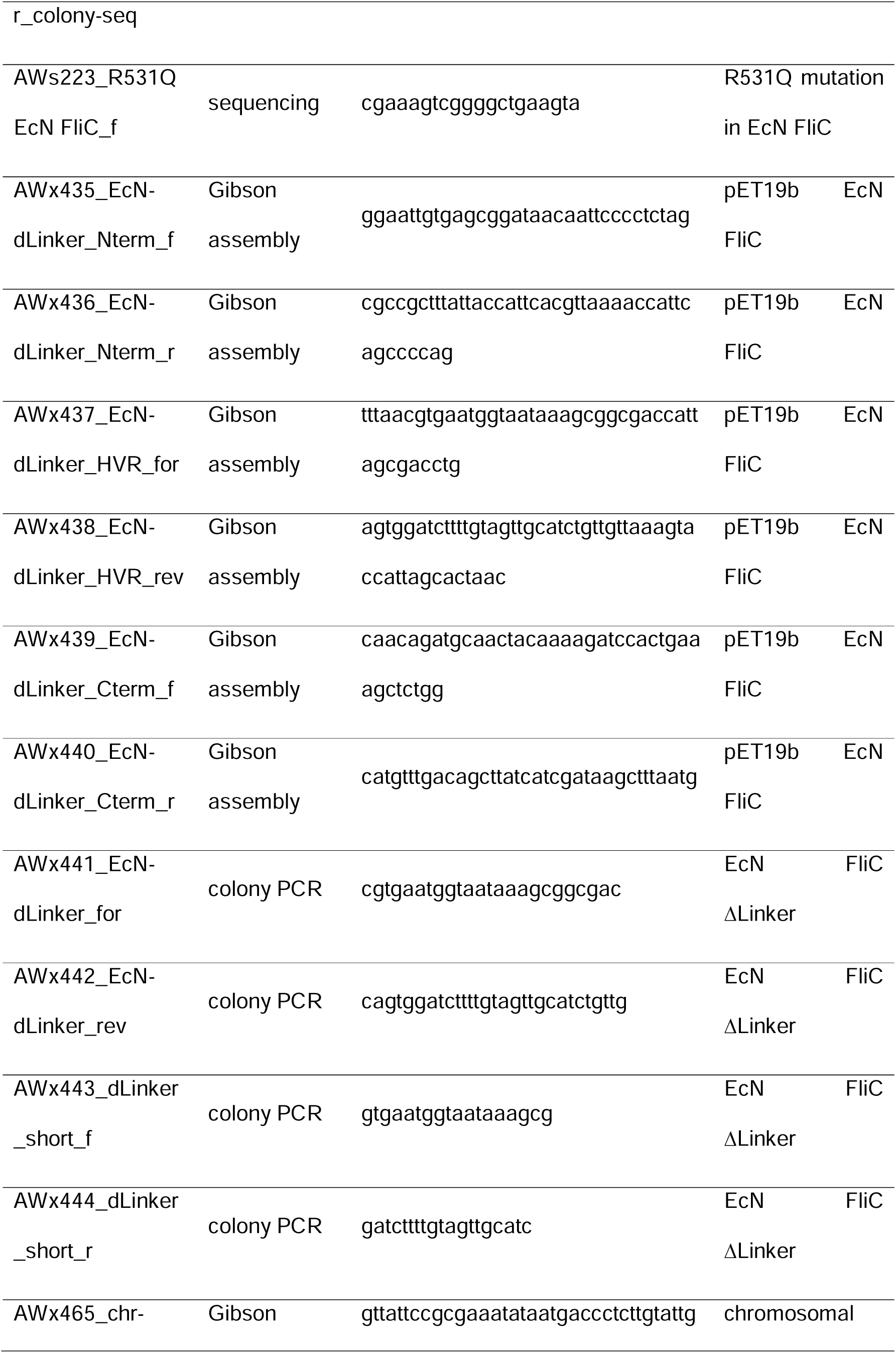
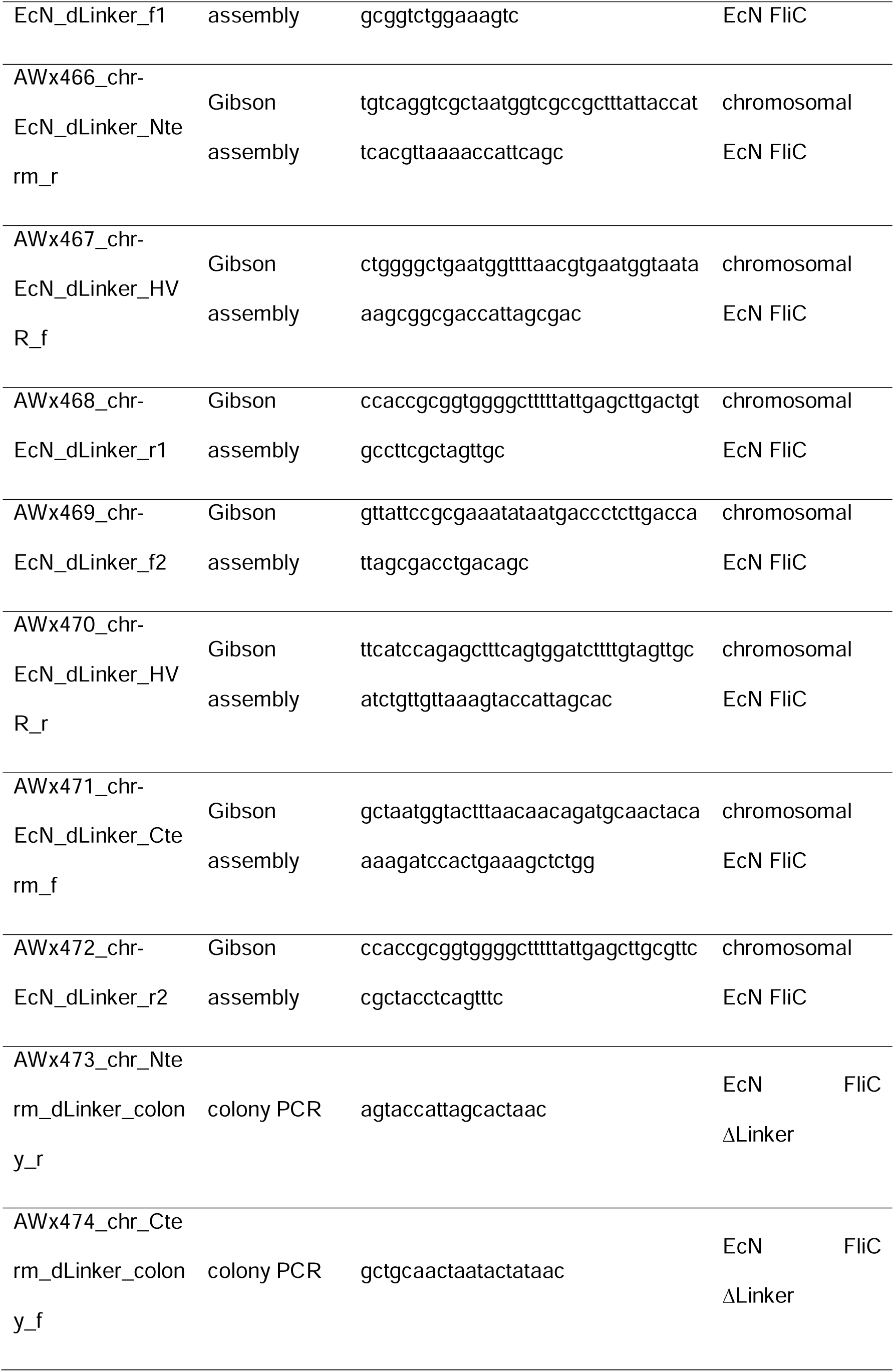
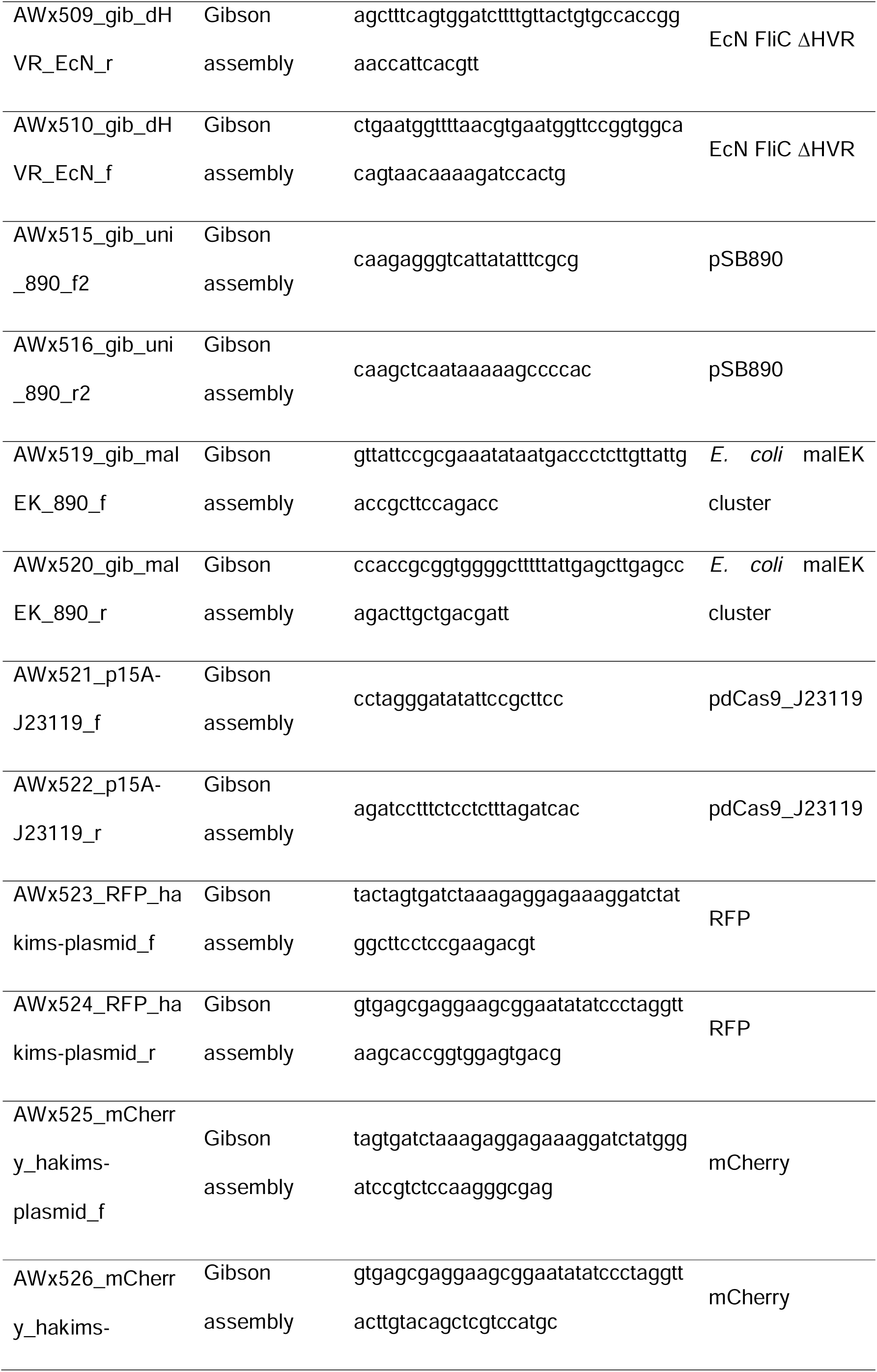
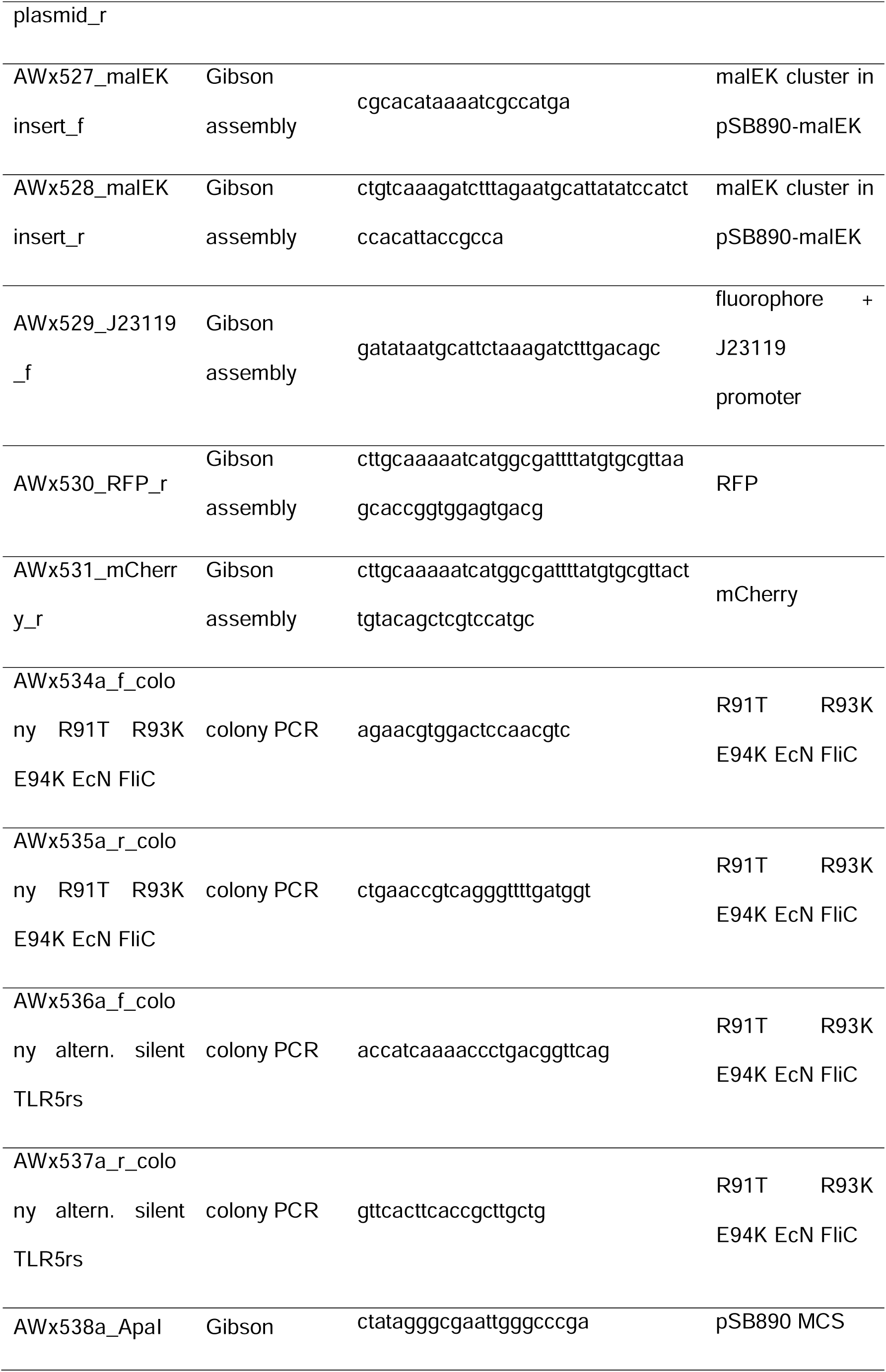
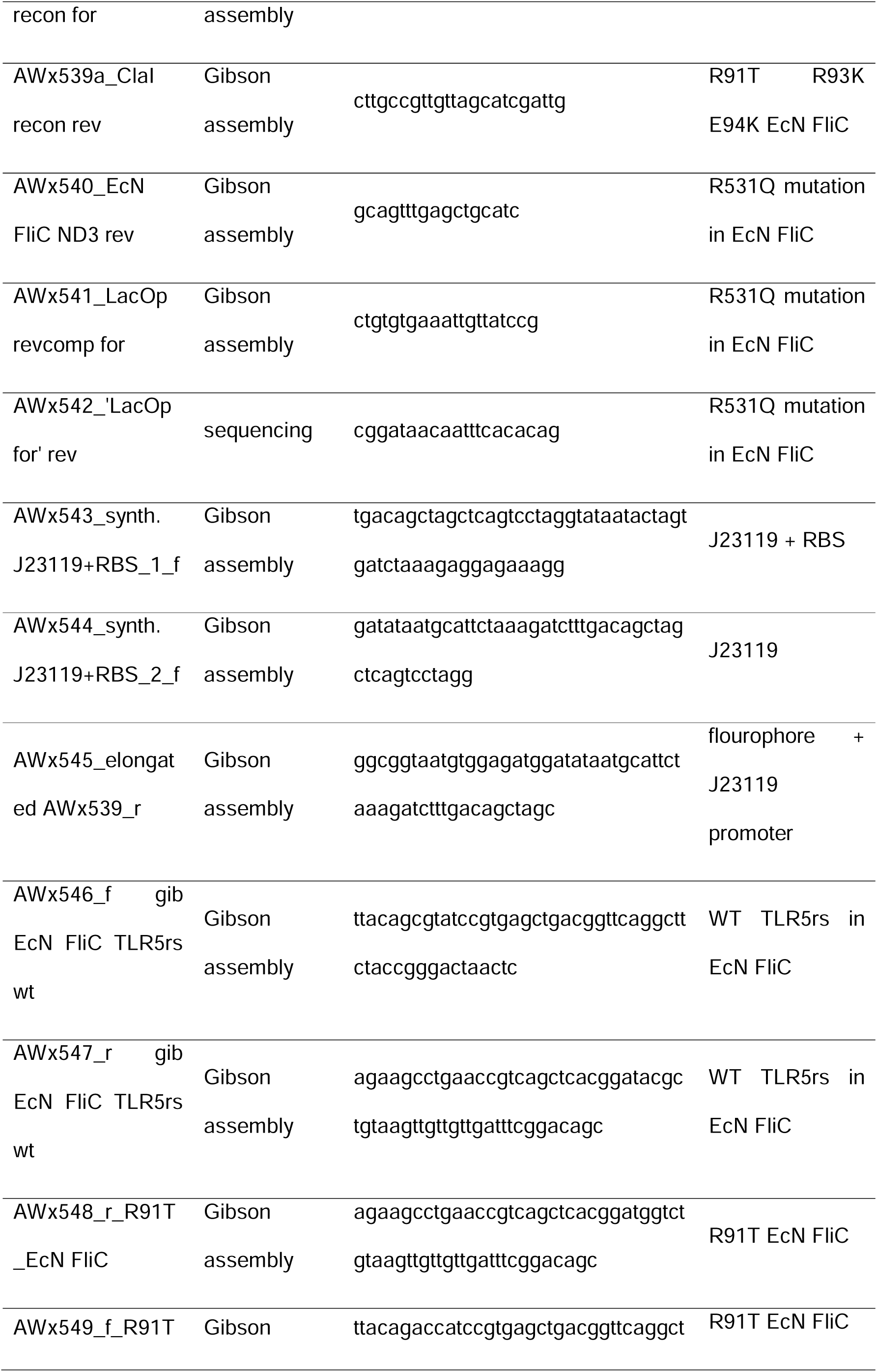
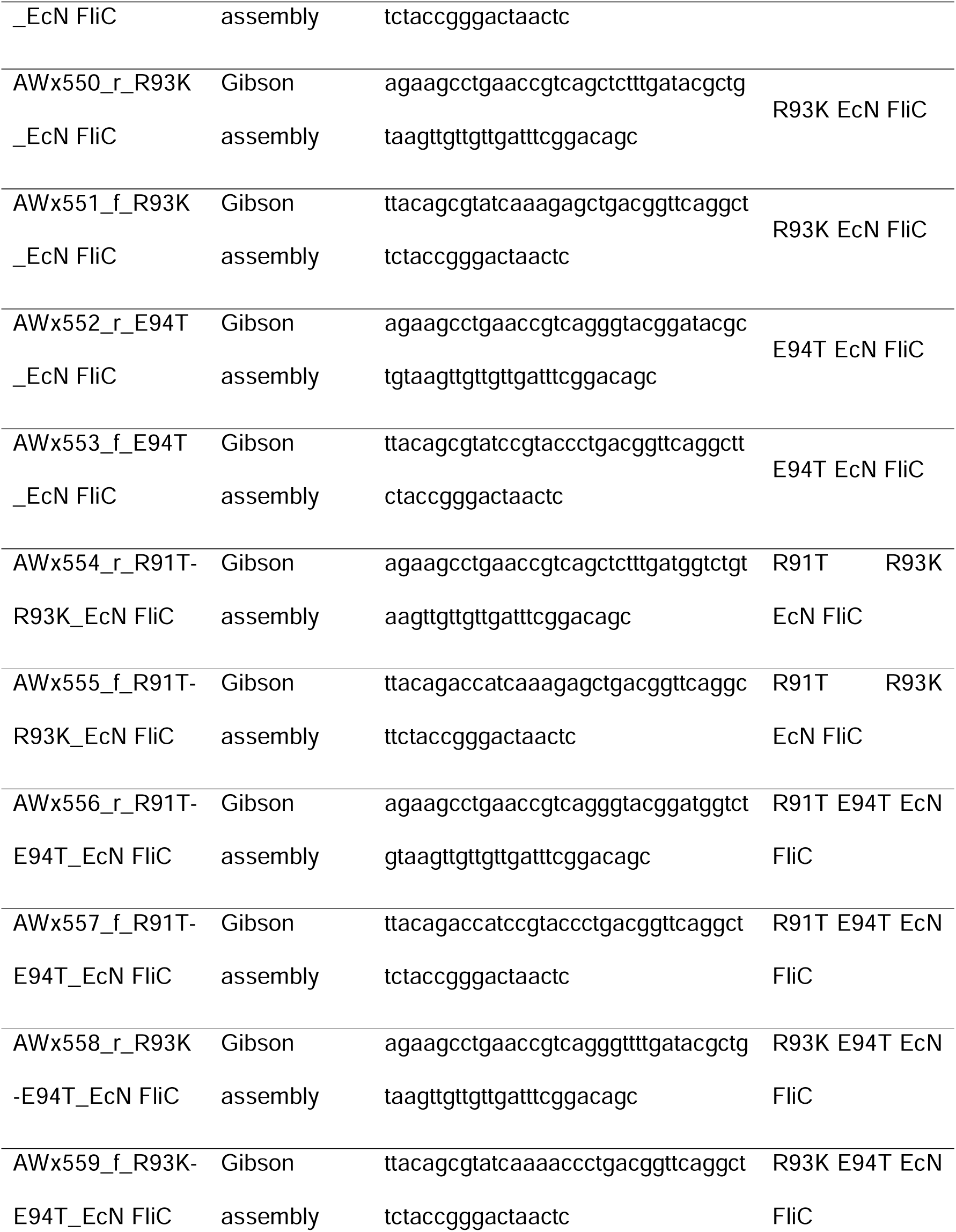
Primers.

**Supplementary table S3:** Plasmids.

| Name | Insert | Backbone | Resistance |
| --- | --- | --- | --- |
| pJJ001 | full EcN FliC + 8xHis-tag TEV cleavable | pET19b | Ampicillin |
| pJJ002 | none | pET19b | Ampicillin |
| pJJ003 | EcN FliC ΔLinker | pET19b | Ampicillin |
| pJJ004 | upstream chromosomal flanking region + N term<br>of EcN FliC lacking N term Linker | pSB890 | Tetracyclin |
| pJJ005 | C term of EcN FliC lacking N term Linker +<br>downstream chromosomal flanking region | pSB890 | Tetracyclin |
| pJJ006 | EcN FliC lacking HVR + chromosomal flanking<br>regions | pSB890 | Tetracyclin |
| pJJ008 | EcN FliC R91T R93K E94T | pGEM-T easy | Ampicillin |
| pJJ007 | intergenomic region + mal E and K cluster<br>beginning | pSB890 | Tetracyclin |
| pJJ009 | EcN FliC R91T R93K E94T R531Q | pGEM-T easy | Ampicillin |
| pJJ010 | malEK cluster containing J23119 mCherry in<br>intergenomic region | pSB890 | Tetracyclin |
| pJJ011 | malEK cluster containing J23119 RFP in<br>intergenomic region | pSB890 | Tetracyclin |
| pJJ012 | EcN FliC WT | pGEM-T easy | Ampicillin |
| pJJ013 | EcN FliC R91T | pGEM-T easy | Ampicillin |
| pJJ014 | EcN FliC R93K | pGEM-T easy | Ampicillin |
| pJJ015 | EcN FliC E94T | pGEM-T easy | Ampicillin |
| pJJ016 | EcN FliC R91T R93K | pGEM-T easy | Ampicillin |
| pJJ017 | EcN FliC R91T E94T | pGEM-T easy | Ampicillin |
| pJJ018 | EcN FliC R93K E94T | pGEM-T easy | Ampicillin |
| pJJ019 | EcN FliC R531Q | pGEM-T easy | Ampicillin |
| pJJ020 | EcN FliC R91T R531Q | pGEM-T easy | Ampicillin |
| pJJ021 | EcN FliC R93K R531Q | pGEM-T easy | Ampicillin |
| pJJ022 | EcN FliC E94T R531Q | pGEM-T easy | Ampicillin |
| pJJ023 | EcN FliC R91T R93K R531Q | pGEM-T easy | Ampicillin |
| pJJ024 | EcN FliC R91T E94T R531Q | pGEM-T easy | Ampicillin |
| pJJ025 | EcN FliC R93K E94T R531Q | pGEM-T easy | Ampicillin |

**Supplementary table S4:** Purification buffers. Buffers used for the purification of FliC. Two different size exclusion buffers were used, for crystallization experiments the SEC-buffer, and for biological experiments DPBS-buffer. The pH of the Buffers where adjusted at 4°C.

| Buffer | Components |
| --- | --- |
| Lysis Buffer / His<sub>A</sub> Buffer | 50 mM Tris pH 8.0<br>300 mM NaCl |
| His<sub>B</sub> Buffer | 50 mM Tris pH 8.0<br>300 mM NaCl<br>500 mM imidazole |
| SEC-Buffer | 20 mM HEPES pH 7.5<br>150 mM NaCl |
| DPBS-Buffer | 0.90 mM CaCl<sub>2</sub><br>0.49 mM MgCl<sub>2</sub><br>2.67 mM KCl<br>1.47 mM KH<sub>2</sub>PO<sub>4</sub><br>137.93 mM NaCl<br>8.06 mM Na<sub>2</sub>HPO<sub>4</sub> |

